# Optimality with room to vary: stiff and sloppy modes in a sensory population

**DOI:** 10.64898/2026.01.30.702802

**Authors:** Kyle Bojanek, Olivier Marre, Stephanie E. Palmer

## Abstract

Populations of sensory neurons are thought to be shaped by selective pressures for optimal information transmission, yet real neural circuits display substantial variability across stimulus repeats, across time, and between individuals. Reconciling this variability with normative theories requires understanding not only the optimal code, but also how performance changes under perturbations of that code. Here we analyze large-scale recordings from retinal ganglion cells responding to diverse naturalistic movies and spatial noise. We fit low-rank Ising models, recast as interpretable latent variable models, revealing that retinal population activity is well described by a small number of collective modes tightly coupled to the visual stimulus. The spatial receptive fields of the leading latent variables closely align with the principal components of masked natural images, consistent with efficient coding. Perturbation analyses based on the Hessian of the efficient coding objective show that this agreement is concentrated along stiff directions, where small deviations strongly degrade performance, whereas sloppy directions tolerate large variability. Across stimulus ensembles, the same stiff latent modes form a stable backbone of population interactions that generalizes between scenes and supports both efficient stimulus reconstruction and predictive coding of future inputs. These results show how sensitivity to perturbations structures sensory population codes, allowing normative optimality to coexist with rich variability.

## INTRODUCTION

Across sensory modalities, biological sensors are confronted with high-dimensional, complex input. Because sensory systems have limited representational capacity, they must devote their responses to the most behaviorally-important features; activity cannot be wasted on irrelevant details. Normative theories seek to formalize which aspects of the input *should* be encoded. Efficient coding theory [1–4] treats all stimulus information as equally valuable and asks how a system should allocate limited resources to represent as much input information as possible. Many systems show strong agreement between efficient coding predictions and their observed encoding properties, including the visual system [3–11], the auditory system [12–14], and the olfactory system [15, 16].

Recent work, however, suggests that not all information is equally useful, and predictive information – about the future state of the input – may be particularly important in visual circuits [17–21]. There is emerging evidence that this principle can describe retinal processing across species [22], primary visual cortex connectivity [23], receptive fields [24], and the transformation of stimulus information up the visual hierarchy [25]. Regardless of whether the objective is efficiency, prediction, or some combination of, these theories posit a single optimal code. Yet, in practice, neural population codes appear highly variable across individuals of the same species [26–30].

Reconciling intraspecific variation with optimality requires going beyond a single best solution and considering the landscape of alternative codes that perform nearly as well. When these alternatives yield similar performance, selective pressure to converge on a single solution is weak, and substantial variability is possible [31]. The extent of this variability depends on robustness: how sensitive performance is to perturbations of the preferred solution. Work in physics [32–35] and machine learning [36–38] shows that high-dimensional systems often con-tain many *sloppy* directions in parameter space, where variation is tolerated, and only a few *stiff* directions, where deviations are costly. Optimality principles are therefore most constraining along stiff dimensions, and their explanatory power declines as sensitivity decreases. If neural population codes share this structure, then we should expect them to match theoretical optima along a small set of stiff features, while tolerating variability along many sloppy directions. Moreover, these key features, themselves, may change in time as stimulus statistics change.

Sensory population codes are highly adaptable across time [39–44]. Natural visual signals evolve on multiple timescales, from rapid shifts in global brightness caused by passing clouds, to slower transitions across day–night cycles and seasons. Movement between environments further alters scene statistics [45]. Efficient coding theory has drawn predictions from natural stimulus statistics across sensory modalities [13, 14, 46], but much less is known about how these statistics change across the diverse contexts organisms must navigate. Characterizing this variability can reveal which aspects of optimal codes remain stable and which must adapt. This form of adaptation goes beyond mechanisms like gain control or rescaling – rather than merely adjusting the strength of existing features, the system may need to encode new stimulus features. Indeed, changing selectivity has been observed in retinal ganglion cells, where the same neurons can encode luminance increases or decreases depending on context [47].

Understanding how adaptation, optimality, and variability coexist in sensory systems requires analysis at both the single-cell and population levels. Population-level analysis is especially important in the retina, where there is a large amount of convergence onto downstream targets [48–51], and behavior is driven by many cells rather than single neurons alone. To probe full sensory populations, we use an interpretable latent variable model of retinal ganglion cell spiking activity from a large recorded population. By analyzing these models, we show how perturbation sensitivity, changing stimulus statistics, and behavioral relevance jointly shape the retinal population code.

We find that retinal responses follow efficient coding predictions most closely along the most sensitive, stiff dimensions identified by perturbation analysis. These dimensions define a small set of collective modes that are highly constrained by natural stimulus statistics and relatively stable across different stimulus conditions. In contrast, along insensitive, sloppy dimensions, the retinal population code departs from the theoretical optimum and exhibits substantial variability. Across stimulus conditions, this pattern is consistent with a stable backbone of retinal ganglion cell interactions, as noted in previous work and found to be useful for rapidly inferring which type of natural environment the animal is in [52]. Finally, when evaluated on a predictive information task, these encodings perform well under both efficiency and prediction objectives, consistent with recent machine learning findings [53]. Together, these results provide a framework for understanding how a biological sensory population can be simultaneously near-optimal, adaptable, and variable.

## RESULTS

### A. Retinal population responses to diverse naturalistic and noise stimuli

To test how efficient coding predictions hold across diverse contexts, responses from 93 retinal ganglion cells were recorded during five naturalistic stimuli (videos of bushes, water, leaves, fish, and self-motion through brush) and a checkerboard stimulus (Fig. 1A). The diversity of these contexts tests how efficient coding predictions shift with changing stimulus statistics, while also enabling assessment of coding robustness and behavioral relevance.

**FIG. 1.**
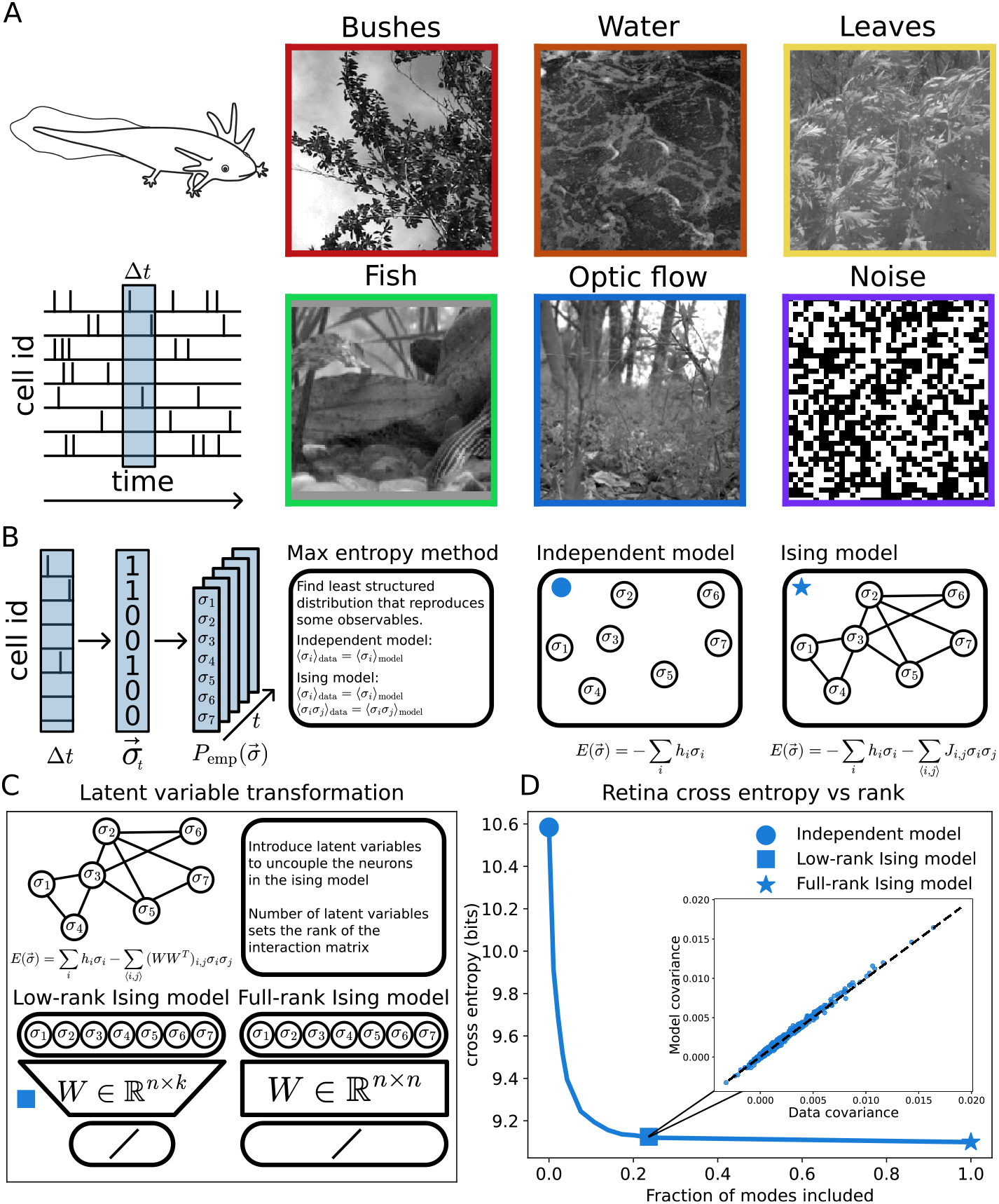
The retinal population code has a compressible structure. **A)** Spiking activity from 93 larval tiger salamander retinal ganglion cells recorded in response to a diverse set of naturalistic and artificial visual stimuli (bushes, water, leaves, fish, optic flow, and checkerboard noise). **B)** The empirical distribution of population spike patterns is modeled using different maximum entropy models, including an independent-neuron model and a full-rank Ising model. Interpolating between these two models is the rank-k Ising model, which constrains the rank of the interaction matrix *J*_*i,j*_ . **C)** The Ising model can be reformulated as a latent-variable model, where the rank of the interaction matrix determines the number of latent variables and the effective dimensionality of population interactions. **D)** Cross entropy of retinal responses to optic flow stimuli as a function of the fraction of interaction modes included in the Ising model. Model performance rapidly saturates with increasing rank. *Inset:* Comparison of pairwise covariances predicted by the low-rank model and measured from data, showing close agreement.

### B. Population activity is well described by a low-rank latent variable model

Efficient coding theory refers to a family of related ideas; here we focus on which stimulus features should be encoded and which ignored. For many combinations of signal and noise statistics, principal components analysis (PCA) provides an optimal ordering of measurements, with successive components capturing progressively less information about the input [54–58]. Under this view, retinal ganglion cells are expected to encode the top principal components of naturalistic stimuli. This possibility was explored previously, but rejected because the resulting spatial filters did not resemble classical retinal receptive fields [59]: the top principal components of natural images are broad and global, whereas retinal receptive fields are spatially localized.

However, retinal ganglion have a rich correlation structure and encode information collectively [60–67]. As a results, the relevant “features” of the code many not be evident at the level of single neurons, but only at the level of the full population activity. This motivates re-examining efficient coding predictions in terms of population-level modes rather than individual receptive fields.

Previous work has examined the population structure of the retina using pairwise maximum-entropy (Ising) models, the least structured model consistent with observed firing rates and pairwise correlations. These models describe the probability of a population response 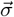 as

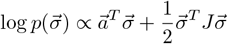

where 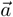 controls individual neuron firing probabilities and the interaction matrix *J* captures pairwise correla-tions. While full-rank Ising models can fit retinal ganglion cell activity well, they place no constraints on the complexity of *J* (Fig. 1B). In contrast, recent work suggests that retinal population activity has a compressible, low-dimensional structure [68, 69], implying that the interaction matrix *J* should also be low-dimensional (Fig. 1C).

We therefore fit low-rank Ising models to the data, in which the interaction matrix is parameterized as *J* = *W W* ^*T*^, where *W* ∈ ℝ^n,k^ and *k* ≪*n*. This explicitly constrains *J* to rank *k* and enforces that correlations arise through a small number of shared latent variables. As the rank, *k*, increases, model performance, assessed by both cross-entropy (Fig. 1D) and reproduction of the empirical correlation structure, saturates rapidly (Fig. 1D). A relatively small number of latent dimensions is sufficient to capture most population statistics, both for naturalistic movies and for checkerboard noise (see supplement figures 1 - 8). This indicates that low dimensionality is an intrinsic property of the retinal population code, not simply induced by correlations in the natural stimuli.

Compared to a full-rank Ising model, the low-rank formulation explains population correlations using far fewer parameters. The latent degrees of freedom can be interpreted as collective variables (see Methods). Rather than interacting directly with each other, retinal ganglion cells interact indirectly through their shared dependence on a small set of latent variables [70, 71] (Fig. 1C). This representation in not just a dimensionality-reduction device, it defines a concrete, probabilistic model of how population-level features are encoded. Studying the coding properties of these collective modes offers a principled way to probe stimulus representations that emerge only at the population level, and connect them to efficient coding principles.

### C. Latent variables have well-defined receptive fields and are strongly stimulus driven

To understand how the latent variables relate to the visual input, we first need to fix a particular representation of the latent space. In the low-rank Ising model, the interaction matrix satisfies *J* = *WW* ^*T*^ . This factorization is invariant under orthogonal transformations of *W*, i.e., replacing *W* with *W U* for any orthogonal matrix *U* leaves the model unchanged because *J* = *WUU*^*T*^ *W* ^*T*^ = *WW* ^*T*^ . Different choices of *U* therefore correspond to different but equivalent organizations of the same latent structure, each emphasizing different combinations of the underlying collective modes.

We choose a representation that is convenient for analysis and interpretation. A natural choice is a variance-maximizing transformation that diagonalizes the covariance of the latent variables, and orders them by variance (Fig. 2A). In this basis, the latent variables are uncorrelated and ranked by how much shared variability in the population they explain, in close analogy to principal components. However, unlike standard PCA, these latent variables are defined for nonlinear, binary neural responses rather than Gaussian signals, so they can deviate substantially from classical principal components [72, 73]. We verified that alternative orthogonal transformations produce qualitatively similar conclusions (Supplementary Information).

**FIG. 2.**
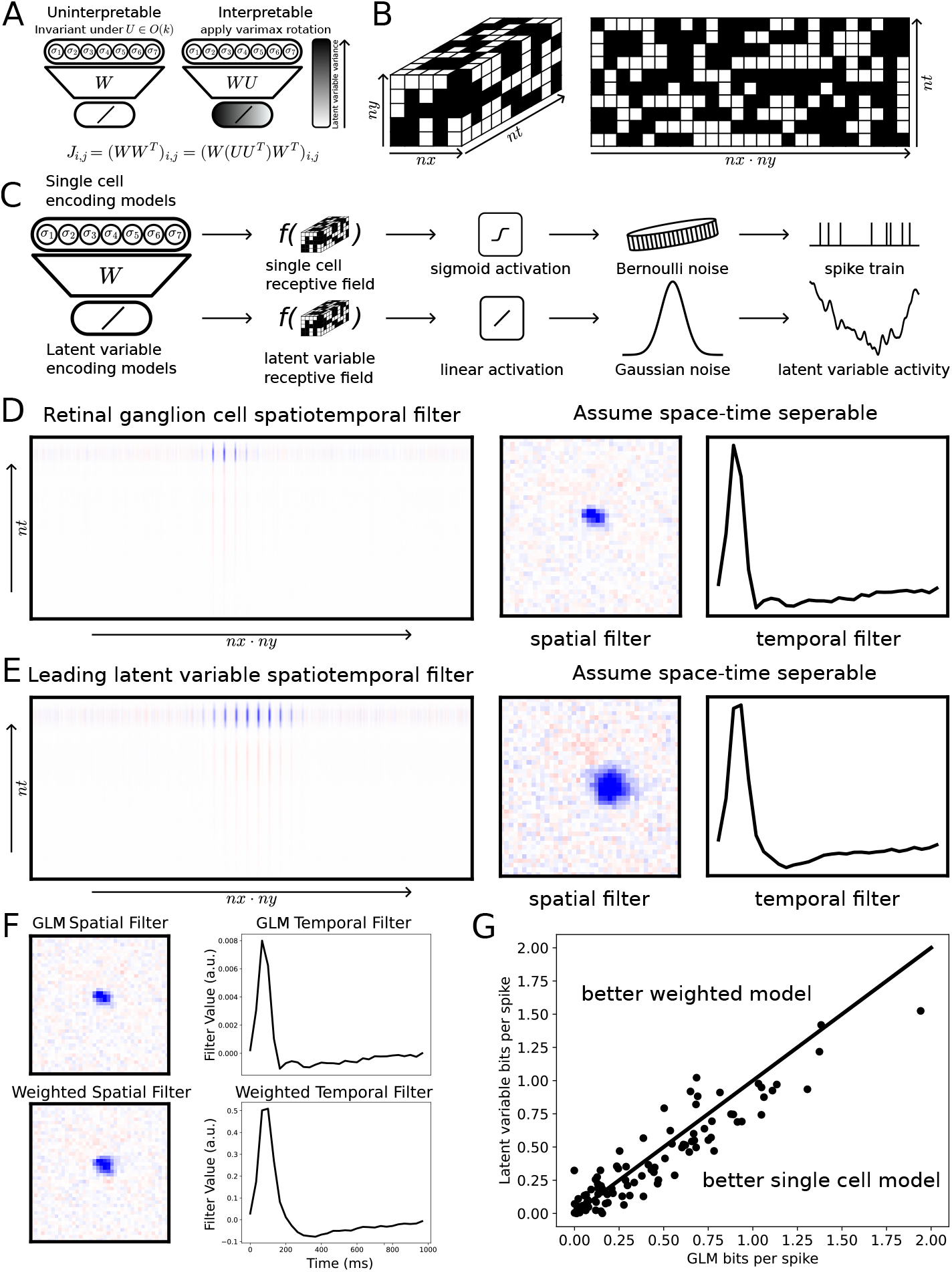
Low-rank Ising model latent variables are sensitive to the stimulus. **A)** The latent-variable representation of the Ising model is invariant under orthogonal transformations of the weight matrix *W* (*UU* ^*T*^ = *I*). An orthogonal matrix that diagonalizes the latent covariance matrix and orders latent variables by explained variance. **B)** Receptive field properties of latent variables are characterized using the checkerboard noise stimulus. **C)** Encoding models for individual retinal ganglion cells are fit using Bernoulli generalized linear models (GLMs) to predict spike probabilities, latent variable responses are modeled using linear regression. **D)** Inferred spatiotemporal filters for individual retinal ganglion cells are well-approximated as space– time separable, allowing decomposition into the outer product of a spatial and a temporal filter. An example cell is shown. **E)** Inferred spatiotemporal filters for latent variables are also space–time separable. Filters for the leading (highest-variance) latent variable are shown. **F)** Single-cell receptive fields inferred directly from GLMs are compared with receptive fields constructed as weighted averages of latent-variable receptive fields, using weights given by the matrix *W* . **G)** Encoding performance of the weighted receptive-field model and independently fit single-cell GLMs, measured in bits per spike on held-out test data for each cell. The two models achieve comparable performance.

With this variance-ordered latent basis fixed, the stimulus dependence of both retinal ganglion cells and latent variables can be examined using a checkerboard noise stimulus (Fig. 2B). Spikes were binned at the stimulus refresh rate, and activity at time *t* was predicted from the stimulus over a temporal window from time *t* to time *t* −*k*. We used logistic regression to model the probability of spiking from the recent stimulus history. For the real-valued latent variables, we used linear regression to predict their activity from the same stimulus window (Fig. 2C).

Consistent with previous work in larval tiger salamander retina, single-cell receptive fields inferred from the checkerboard stimulus were predominantly OFF-center with fast temporal filters. These spatiotemporal filters are well approximated as separable in space and time, and can be decomposed into the outer product of a spatial and a temporal component (Fig. 2D). Strikingly, the leading latent variable, the one with the highest-variance, shows the same qualitative structure. It displays an OFF-center spatial receptive field, a fast temporal kernel, and is space–time separable (Fig. 2E; see Supplementary Information). Thus, the most prominent collective mode of the population inherits familiar single-cell-like tuning, but at the level of shared activity across many neurons.

Because both retinal ganglion cell sand latent variables respond robustly to the stimulus, we can ask how single-cell receptive fields are composed of these collective modes. To do this, we construct a “weighted-latent” model in which the receptive field of cell *i* is expressed as a linear combination of latent-variable receptive fields, with weights given by the coupling matrix *W* . This yields a reconstructed receptive field for each neuron by mapping latent filers back into individual cells with *W* ^*T*^ . These reconstructed receptive fields closely resemble those obtained from independently fit singlecell models (Fig. 2F) and achieve comparable encoding performance when predicting spikes from the stimulus (Fig. 2G).

Together, these results show that retinal encoding is organized around a small set of collective variables that are tightly coupled to the visual stimulus and have welldefined and interpretable receptive fields. Single-cell receptive fields can be understood as specific linear combinations of these population-level modes, indicating that the latent variables provide a compact, stimulus-driven basis for the retinal population code.

### D. Latent variables align with principal components of natural scenes

Single-cell receptive fields of retinal ganglion cells are well characterized, but far less is known about the stimulus tuning of population-level features, modulo some notable earlier papers [65, 74–76]. In the variance-ordered basis defined above, the leading latent variables capture the largest sources of shared variability in the population response. Under an efficient population code, these dominant variables should reflect the dominant structure of the visual input. In particular, we expect them to align with the principal components of natural images, which optimally capture stimulus variance under broad conditions [54–58].

To characterize the structure of natural scenes, principal components were computed from a large natural image dataset (CIFAR-10) [77]. Images were masked to match the region driving the retinal response, and we diagonalized the resulting covariance matrix. As in previous work [58], the leading principal components encode coarse, low-spatial-frequency structure over the masked region, while higher-order components capture progressively finer spatial spatial detail (Fig. 3A).

**FIG. 3.**
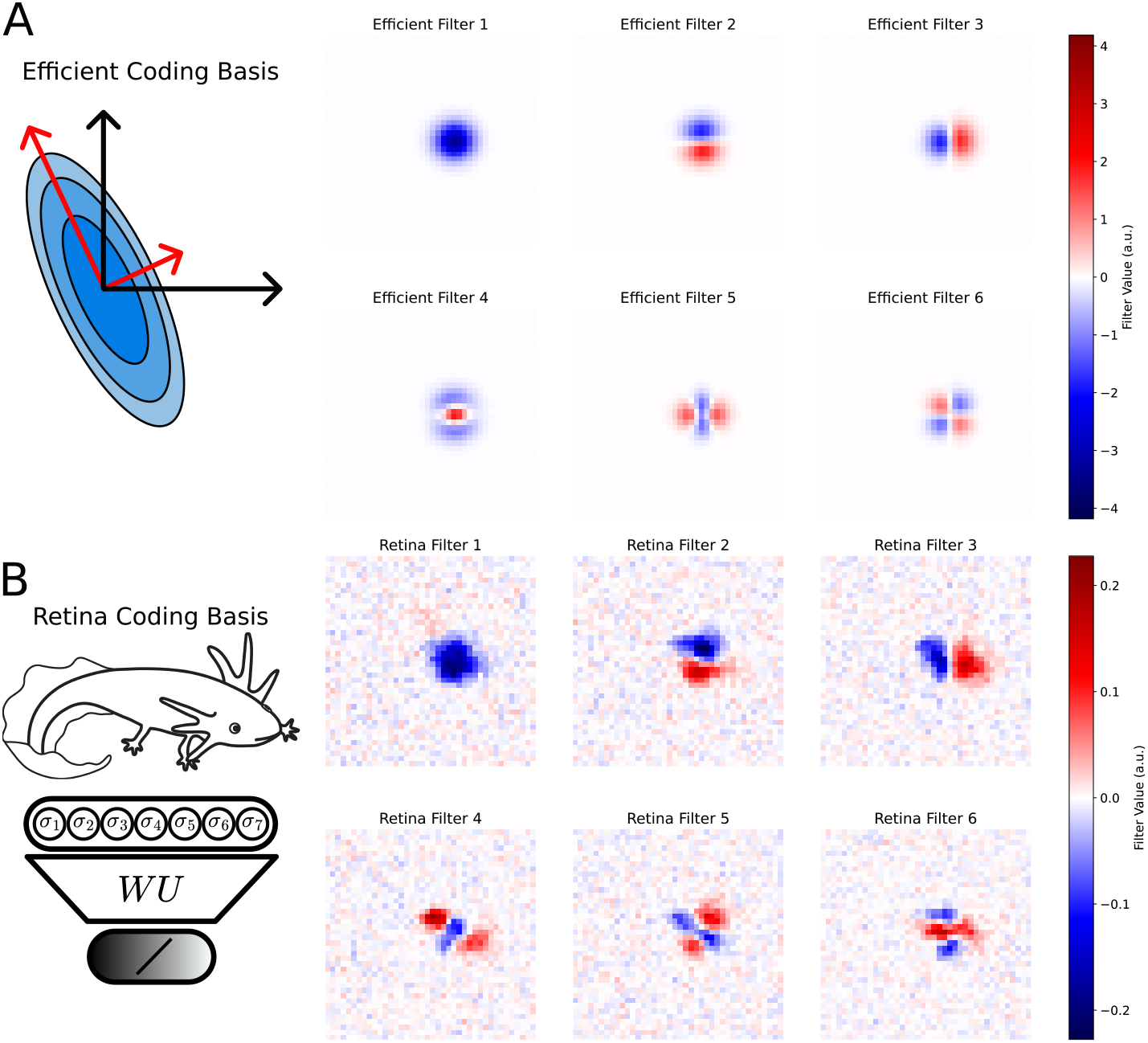
Latent-variable receptive fields are consistent with efficient coding predictions. **A)** Efficient coding basis functions derived from natural scene statistics. Shown are the six most informative spatial filters, corresponding to the leading principal components of a dataset of masked natural images. **B)** Spatial receptive fields of the six highest-variance latent variables inferred from the low-rank Ising model. The inferred latent-variable filters show strong qualitative agreement with the efficient coding basis.

We then inferred spatial receptive fields for the latent variables using the checkerboard stimulus (Methods). When the latent variables were ordered by variance in the retina data, their spatial receptive fields exhibited a strikingly similar hierarchy: the leading latent variables encode coarse low-frequency spatial structure over the retinal field of view, while lower-variance latent variables represent progressively finer spatial features (Fig. 3B). The correspondence between natural image principal components and latent receptive fields is particularly strong for the first three modes.

Importantly, the latent receptive fields were inferred using spatiotemporally uncorrelated checkerboard noise rather than natural images. The observed agreement reflects intrinsic structure in the retinal population code, rather than being simply induced by the correlations in natural stimuli. Together with the low-rank analyses above, these results indicate that the dominant collective modes of the retinal population encode the dominant modes of variation in natural scenes, as predicted by efficient coding theory, but expressed at the level of population activity rather than individual neurons.

### E. Efficient coding predictions are strongest along stiff, perturbation-sensitive directions

The close match between latent receptive fields and natural image principal components suggests strong selective pressure acting on the retina. However, the strength of this selective pressure is not necessarily uniform across inferred retina filters. Some aspects of the code may be tightly constrained because small deviations cause large performance losses (“stiff” directions), whereas others may tolerate large variability with little cost (“sloppy” directions). To understand where efficient coding should most strongly constrain retinal population structure, we analyzed the local geometry of the efficient coding problem.

We considered a simplified version of efficient coding in which the goal is to find a linear projection *x* (a spatial filter) that maximizes stimulus variance *x*^*T*^ *Sx*, where *S* is the image covariance matrix, subject to |*x*| = 1. This is the standard PCA objective and yields the first principal component as the optimal solution. Introducing a Lagrange multiplier *λ* for the unit-norm constraint gives the objective function

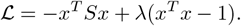

The gradient,Δℒ = 2*Sx* −2*λx*, vanishes at *Sx* = *λx*, so stationary points are eigenvectors of *S*. The second derivative (Hessian) of ℒ, Δ^2^ ℒ = 2*S* 2*λI*, at such a solution has eigenvectors aligned with the principal components of *S*, with eigenvalues 2(*λ*_1_ −*λ*_*i*_), where *λ*_1_ is the variance captured by the leading principal component and *λ*_*i*_ are the remaining eigenvalues. Thus, perturbations that mix the leading component with low-variance components (large *λ*_1_ −*λ*_*i*_) lie along high-curvature, stiff directions where performance degrades rapidly. Perturbations that mix it with other high-variance components (small *λ*_1_ −*λ*_*i*_) lie along flat, sloppy directions where performance is relatively insensitive (Fig. 4A).

**FIG. 4.**
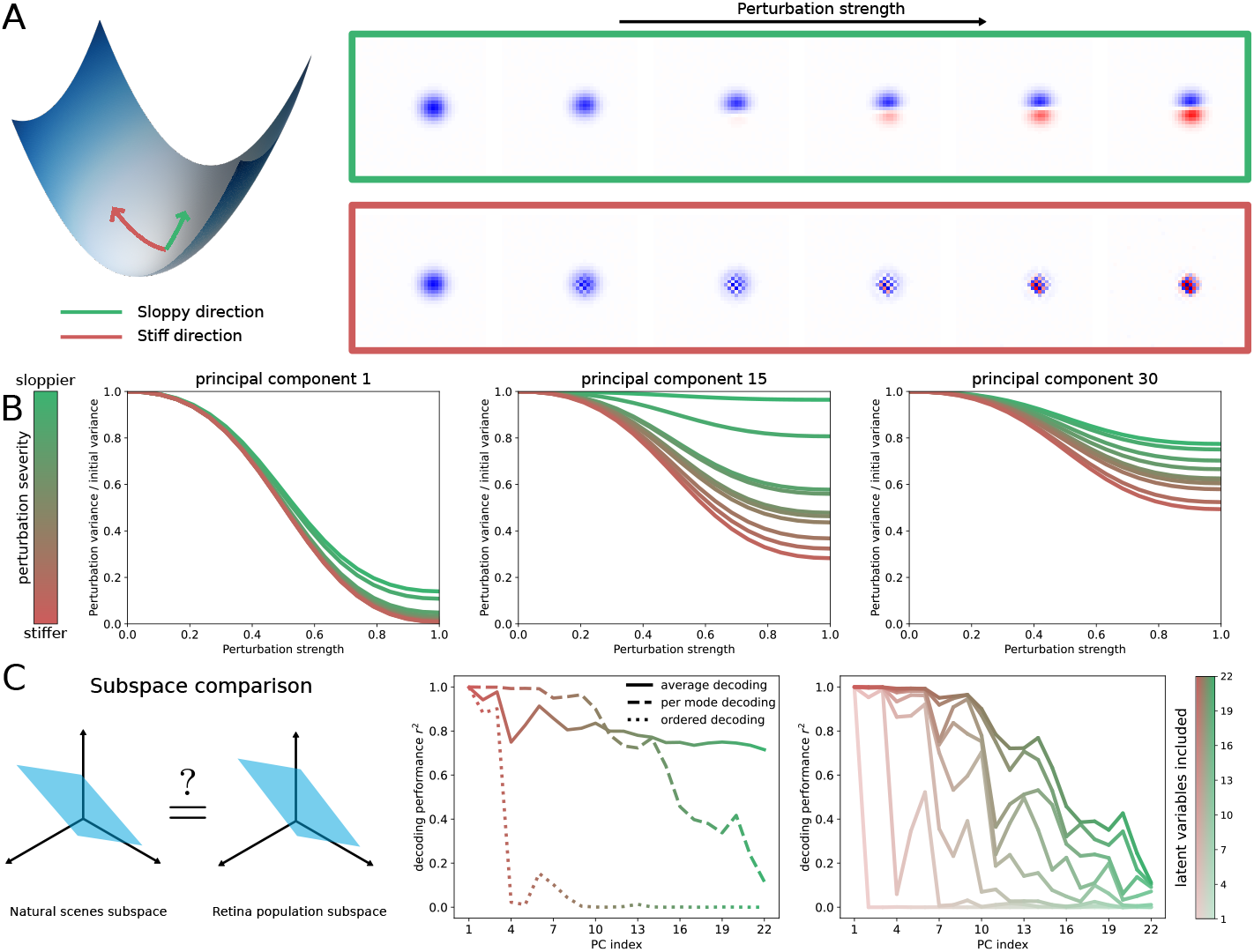
Agreement with efficient coding theory depends on the sensitivity of optimal solutions to perturbations. The loss landscape of the efficient coding objective contains both sensitive (stiff) and insensitive (sloppy) directions. Perturbation of the first principal component towards the second principal component causes the least degradation in performance. Perturbation towards the last principal component causes the most degradation. **B)** Sensitivity to perturbations across multiple principal components. Green curves correspond to perturbations along sloppy directions, while red curves correspond to perturbations along stiff directions. Higher-order principal components exhibit an increasing number of sloppy directions and a reduced relative sensitivity to perturbations. **C)** Comparison of the subspace spanned by retinal latent-variable spatial filters and the subspace spanned by efficient coding spatial filters. Subspace similarity is quantified using three decoding strategies: an average decoder (mean *r*^2^ obtained by using retinal filters 1 through *i* to decode principal components 1 through *i*), an ordered decoder (decoding principal component *i* using retinal filter *i*), and a per-mode decoder (decoding principal component *i* using all 22 retinal filters). The per-mode decoder is further decomposed by restricting decoding to the first *k* retinal filters.

Extending this analysis to the *i*th principal component requires first “deflating” the covariance by removing the contributions of previously learned components:

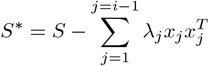

Repeating the Hessian analysis on *S*^∗^ shows that as one moves to higher-index components, the largest curvature eigenvalues decrease roughly in proportion to the eigen-values of the covariance matrix. In other words, the leading components are strongly constrained by the objective, while later components become progressively less sensitive to perturbations. A similar analysis, with similar results, can be done for learning *k* components simultaneously [78, 79]. For masked natural images, this manifests as strong sensitivity of the first principal component to even the least harmful perturbations, and much weaker sensitivity for higher components, which can be altered substantially with minimal performance loss (Fig. 4B–C). This pattern is a direct consequence of the power-law spectrum of natural images [80].

These geometric considerations lead to a concrete prediction: agreement between efficient coding theory and the retinal collective modes should be strongest for the first few components and should systematically weaken with component index. Features corresponding to high-curvature (stiff) directions in the efficient coding landscape should be most conserved across individuals and stimulus conditions, while low-curvature (sloppy) directions should allow substantial variability with little impact on performance.

To test this prediction, we compared the spatial filters of the retinal latent variables to principal components of masked natural images. We projected images onto the subspaces defined by both the PCA filters and the retinal spatial filters, and then evaluated three decoding strategies (Fig. 4C). An ordered decoder enforced a strict one-to-one correspondence: retinal filter *i* was used to decode principal component *i*. Despite this stringent test, the first three retinal filters decoded the first three principal components with high *r*^2^. A more permissive per-mode decoder allowed any combination of retinal filters to decode each principal component, probing overall subspace similarity. Here, good agreement extended to more components, but *r*^2^ declined with component index, as predicted by the sensitivity analysis. Finally, an average decoder used the first *k* retinal filters to jointly decode the first *k* principal components, summarizing performance as an average *r*^2^ over components. This measure likewise decreased as additional, less constrained components were included.

The Hessian analysis and decoding results show that efficient coding provides its strongest, most precise predictions for a small number of stiff, high-variance collective modes. Beyond these modes, the loss landscape becomes increasingly flat, allowing many alternative retinal population codes to coexist with only minor performance differences.

### F. Stable and adaptable components of the population code across visual contexts

The analyses above used principal components computed from a large dataset of natural images, pooled over different image classes (CIFAR-10) [77]. In reality, organisms encounter many distinct visual contexts, each with its own statistics [45]. If the retina is approximately efficient for each context, then the optimal filters may shift with scene class. At the same time, behavior requires stability. Some components of the code should remain useful across contexts. Here we use natural image statistics and retinal data to distinguish which aspects of the retina population code are stable, and which are adaptable.

First, we quantified how the efficient coding solutions, itself, varies across visual environments. Instead of computing principal components on CIFAR-100 pooling over all 100 distinct classes, we computed them separately for each class. For every pair of classes, we compared the subspaces spanned by the principal components using an ordered decoder (as in Fig. 4C). This measures how well component *i* from one class can be reconstructed from component *i* of another (Fig. 5A,B). We found that the subspace similarity declined with the component index (Fig. 5A,B), measured both by the median and maximum. The leading components were highly conserved across classes, while higher components diverged more strongly. Notably, there is a significant drop in similarity at the second component, followed by a return to a nearly identical subspace in the third component. This reflects the fact that components 2 and 3 encode horizontal and vertical features, which share similar variance across classes but can swap order.

**FIG. 5.**
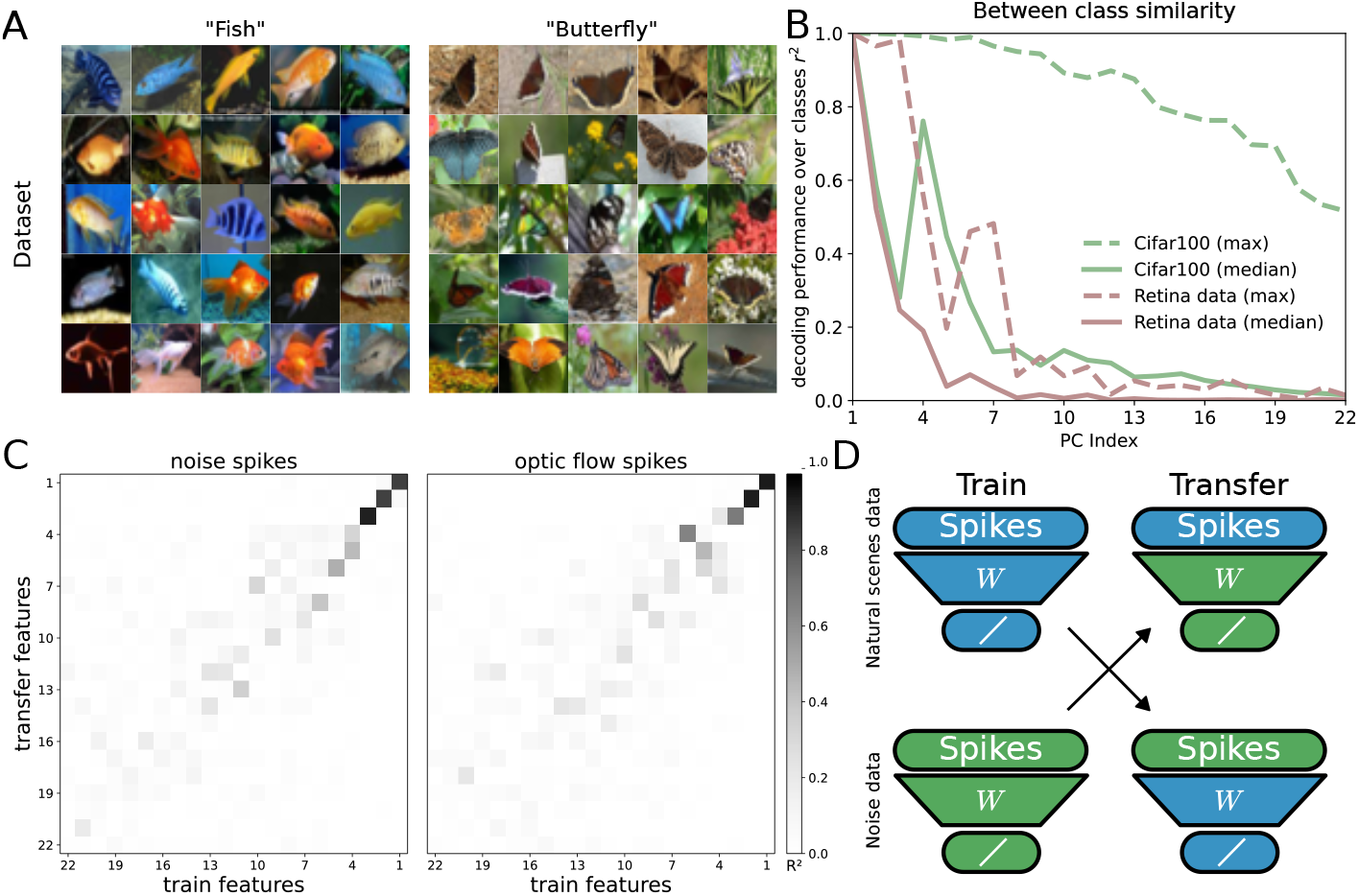
Efficient coding filters change with stimulus statistics) **A)** Organisms exist in a wide variety of contexts and within these contexts natural scene statistics can be very different. Example classes from natural images dataset are shown. For the natural image dataset, the optimal subspace from one class is used to decode the optimal subspace for another class using an ordered decoder (index *i* of subspace 1 decoding index *i* of subspace 2; green). The same analysis is performed for the retina latent-variable subspace (brown). Median and maximum decoding performance are pooled across all 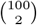 class pairs for efficient coding subspace and across all classes for retina subspace. In both cases, decoding performance decreases with index. **C)** Two stimulus conditions are taken (optic flow and noise) and the latent variable activity from projecting one stimulus conditions spikes onto its own latent variables are compared with the latent variable activity from projecting those spikes onto the other stimulus conditions latent variables. The squared correlation is shown for each pairwise comparison. The top three latent variables are stable across the noise and optic flow stimulus conditions. **D)** A cartoon illustrates the transfer learning procedure.

We then compared the retinal spatial filters to the class-specific efficient filters in the same way. The distribution of ordered-decoder performance across all image classes again showed high similarity for the first few components, followed by a systematic decline with component index (Fig. 5A,B). Thus, the features of the retinal code that best match efficient coding are precisely those that are most stable across natural scene classes. Higher-order components, which are less constrained by image statistics, show more divergence between the retina and any single efficient solution.

Next, we directly probed stability and flexibility in the retinal population code across different stimulus ensembles presented to the same retina. This is tested with a transfer learning experiment. Spikes from one stimulus condition are taken and projected onto the latent space inferred from that condition (“within-condition”), as well as onto latent spaces inferred from other conditions (“across-condition”). We then quantified how well each latent variable from one condition could be predicted from the corresponding latent variable defined in another, using *r*^2^ as a similarity measure (Fig. 5C,D). Across comparisons between the optic flow movie and checkerboard noise, similarity between matched latent variables decreased with latent index: the first few latent variables were strongly conserved, while later ones differed markedly between conditions. Similar trends were observed across other pairs of naturalistic stimuli (Supplementary Information). While the diversity of natural scenes considered places our study outside of the regime where direct perturbative analysis is possible [47, 81], our finding of a stable core of low frequency features agree with previous work highlighting the insensitivity of low-frequency features to perturbation [81].

The natural image analysis and the cross-condition latent comparisons support a common picture. A small set of leading collective modes, those corresponding to the most constrained, high-variance directions in natural scene statistics, form a stable backbone of the retinal population code that generalizes across diverse visual environments. In contrast, lower-variance, sloppy modes are more stimulus dependent and provide an substrate for context-dependent adaptation.

### G. A shared backbone supports both efficient representation and prediction

So far, we have focused efficient coding, which assumes that all information is equally valuable. For moving ob-servers in moving environments, information that predicts future sensory input may be especially important. This problem is acute in the retina, where phototransduction and synaptic integration introduce significant lags that must be overcome by active prediction. We ask whether the same population-level features that support efficient encoding of the current visual input also support accurate prediction of future inputs.

To compare efficient and predictive objectives, we constructed two set of idealized spatial features from natural movies. The efficient coding features were defined as the top principal components of the covariance matrix, *C*_*t*_, capturing directions that best reconstruct the current visual input frame from its low-dimensional projection. The predictive features were defined using reduced rank regression, between frames at time *t* and at a later time *t* + *k*, capturing directions that optimal predict future visual inputs from the present frame. As a baseline, we also considered random orthogonal bases, and we compared all of these to the spatial features derived from the retinal latent variables (the “retina basis”) (Fig. 6A).

**FIG. 6.**
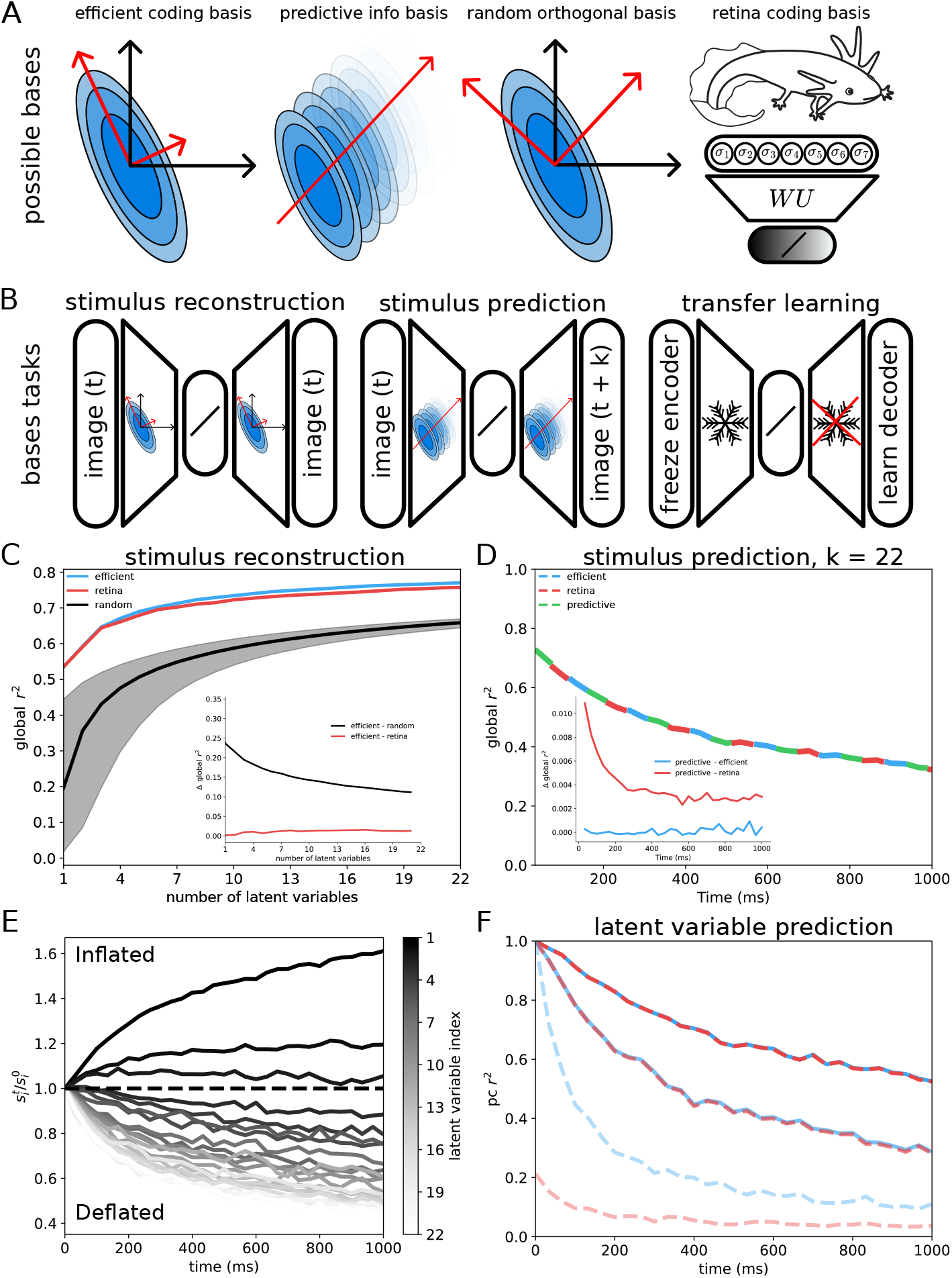
Predictive features closely resemble efficient coding features. **A)** Schematic of the filter bases under comparison: efficient coding, predictive information, random orthogonal, and retinal. **B)** Task definitions. For stimulus reconstruction, masked natural images at time *t* are projected into a low-dimensional latent space and decoded to reconstruct the same image. For stimulus prediction, masked frames at time *t* are projected into a latent space and used to predict the frame at time *t* + *k*. For transfer learning, an encoder learned on one task is frozen and evaluated on a different decoding task. **C)** Stimulus reconstruction performance, measured by global *r*^2^ across pixels on held-out data, as a function of the number of latent variables for efficient, retinal, and random orthogonal bases. Random orthogonal bases shown with median, and fifth and ninety-fifth percentile error bars. **D)** Stimulus prediction performance (*k* = 22) measured by global *r*^2^ as a function of time. Efficient, retinal, and predictive bases achieve similar prediction accuracy. Inset: pairwise performance gaps (difference in global *r*^2^). **E)** Relative change of singular values of the predicted future, normalized by the corresponding equal-time singular values. As prediction horizon increases, the leading three singular mode captures an increasing fraction of the variance at the expense of higher-order modes. **F)** Prediction performance, measured by *r*^2^ with different natural image principal components, as a function of time. Curves are shown for components 1, 2, and 22 (all others in supplement); opacity decreases with component index.

We evaluated each feature set using a transfer-learning scheme that separates encoding from decoding (Fig. 6B). For each basis, we kept the encoder fixed (projection of images onto the basis) and trained a new linear decoder for each task. This allowed us to ask how well a given set of features (optimized for one objective) could support other objectives when only the readout is adapted.

On a stimulus reconstruction task (reconstructing the current frame), the efficient coding basis was, by construction, optimal. Remarkably, the retinal bases achieved nearly identical performance, as measured by global *r*^2^ across pixels (Fig. 6C). The agreement was especially strong for the first three latent dimensions and declined only slightly for subsequent dimensions, (Fig. 6C, inset), consistent with the stiff-sloppy structure inferred from the Hessian analysis. The performance gap between each basis and the efficient coding optimum was small for the retina basis and much larger for random features.

We next turned to a stimulus prediction task, in which frames at time *t* are used to predict frames at time *t*+*k* for a range of temporal offsets. The predictive features were, by construction, optimal for this task. When we evaluated the retina and efficient coding bases in the same transfer-learning framework (fixed encoder, learned decoder), we found that all three bases (efficient, predictive, and retinal) achieved comparable prediction accuracy (Fig. 6D) as measured by global *r*^2^ across pixels. Again, the gap relative to the predictive optimum was small for both the efficient and retinal features and much larger for the random features (Fig. 6D, inset). Notably, the performance gap for the retina basis remained small at short times but decreased modestly at longer prediction horizons.

To understand why these distinct objectives lead to similar performance, we analyzed the structure of the stimulus covariance as well as the reduced rank regression predictions. Projecting natural movies onto the efficient coding basis reveals that the same eigenvectors that diagonalize the covariance matrix, *C*_*ij*_, also approximately diagonalize the past-future cross-covariance 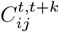, across a broad range of *k* (Supplementary Information). This agreement is strongest for the leading modes, implying that efficient and predictive coding share a common set of dominant features. Although the eigenvectors are similar, their relative importance changes over time. The singular values of the reduced rank regression predictions, normalized by their equal-time values, show that the leading three modes grows in relative importance as the prediction horizon increases, while all higher-order modes decay (Fig. 6E). Consequently, prediction at long times is dominated by the first three component, which is the most accurately captured by the retinal bases. This explains why the performance gap between predictive and retinal features shrinks with increasing prediction horizon.

This effect is also evident when decoding individual natural image principal components over time. Prediction performance decays most slowly for the first principal component in both the efficient coding and retinal bases, whereas higher-order components decay more rapidly and are represented less accurately by the retina (Fig. 6F). Low spatial frequency components therefore dominate long-term prediction, consistent with the coupling between spatial and temporal frequencies in natural scenes [82].

Together, these results show that the same small set of leading collective modes (identified independently of measurements of retinal activity) forms a shared backbone that supports both efficient representation of current inputs and accurate prediction of future inputs. This analysis does not determine if the retina is more suited to reconstruct stimuli or predict future input, rather it suggests that a similar basis is approximately optimal for both. The main change in structure between the optimal representation for the efficient coding problem and the prediction problem, is not what features are relevant, but how well measured each feature is. For prediction noise needs to be more suppressed in lower-frequency modes and is more permissible in higher-frequency modes. Noise correlations may play a particularly important roll in controlling the frequency dependent signal-to-noise ratio, as they have been shown to have frequency dependent effects on information transmission [83]. The usefulness of features across multiple tasks aligns with recent machine learning work showing that features learned for one objective (e.g., reconstruction) can be highly transferable to others (e.g., prediction) [53]. In the retina, this shared backbone provides a concrete mechanism by which a single population code can simultaneously satisfy multiple normative objectives while still allowing variability and adaptation along less constrained directions. Future work, could investigate which time horizon populations of retinal ganglion cells are most well suited to predict.

## DISCUSSION

The population code for retinal ganglion cells can be described effectively with a small number of latent variables that are strongly coupled to the stimulus. A stimulus encoding model constructed from latent variables shows comparable performance to single-cell spike triggered averages. While the response properties of single cells are understood, our results shed new light on how the stimulus is collectively encoded by retinal populations. Our analysis, while preliminary, reveals several interesting and interpretable features in this collective code and makes connections to efficient coding. Further work will be needed to fully understand how the response of these collective variables change with changing stimulus statistics.

A central observation is that the most important latent variables are conserved across stimulus conditions. Retinal encoding models often fail to generalize far outside the stimulus ensemble on which they were trained, yet a subset of latent variables forms a stable “core” of population interactions, as suggested previously [52]. Understanding this stable core may provide a route to describing population codes across natural scenes, and to identifying which aspects of the code are under the strongest constraints.

Our results also motivate a re-examination of a classical efficient coding proposal. The naive source-coding idea, that the retina directly measures stimulus principal components, was previously rejected for not giving physiologically realistic receptive fields. Here we show that this objection is resolved at the population level. When we analyze the receptive fields of collective variables, rather than individual cells, we find that encoding the principal components of natural images is in fact reasonable. This suggests that efficient coding at the level of hidden latent variables may give rise to sparse, localized receptive fields in visual spiking neurons. Clarifying this relationship, and understanding how retinal circuitry implements the observed collective modes, will require further work to reveal the mechanistic underpinnings of these effects. The fact that similar latent modes appear under both noise and naturalistic stimuli indicates that they are not simply induced by external correlations, but instead reflect intrinsic structure in retinal circuitry.

Our findings highlight two additional features of natural vision that are relevant for normative theories. First, there is substantial variability in natural scene statistics across stimulus classes, and second, features that are useful for one computational objective often transfer to others. We still find meaningful agreement between neural data and normative theories, but these two observations suggest that the “inverse” use of normative theories may be particularly useful. Rather than only asking whether a given input ensemble and objective function can reproduce the observed retinal code, we can also ask which input statistics would make the measured latent receptive fields optimal[84, 85]. Because the loss landscape contains many flat directions, such inverse problems will generally admit many near-equivalent solutions, and care should be exercised in evaluating them. Nonetheless, this approach may be useful for distinguishing between competing theories of optimality.

The analysis of the Hessian of the efficient coding problem plays a key role in interpreting our results. By characterizing how performance degrades under perturbations of the optimal solution, we showed that the efficient coding landscape contains a small number of stiff directions, where deviations are costly, and many sloppy directions, where variation is tolerated. Our comparison between retinal population codes and efficient coding predictions is consistent with this structure: strong agreement is observed along stiff directions, while greater variability and weaker correspondence appear along sloppy directions. This suggests a general strategy for using normative theories in biological systems. Instead of asking whether a system matches the unique optimum, we should ask where in parameter space deviations are expected to be small, and where variability is likely.

This perspective may be especially useful for understanding variability across species. Efficient coding theory is often used to define a single optimal code, which seems at odds with the diversity observed in nature. However, if real systems are governed by high-dimensional objectives with many flat directions, then variation along those directions is expected. Analyzing the Hessian of an appropriate normative objective can therefore indicate where interspecies variability is likely to be found and where it is unlikely [31].

## ACKNOWLEDGMENTS

KB would like to thank Cheyne Weis for useful discussions. This work supported by the Polymaths Program from Schmidt Sciences, LLC; the Physics Frontier Center for Living Systems through the National Science Foundation award NSF PHY-2317138; the NSF-Simons National Institute for Theory and Mathematics in Biology, awards NSF DMS-2235451 and Simons Foundation MP-TMPS-00005320; the University of Chicago Materials Research Science and Engineering Center, award NSF DMR-2011854; the Center for the Physics of Biological Function, NSF PHY-1734030.

## MATERIALS AND METHODS

### A. Multielectrode array recordings from larval salamander retina

Voltage traces were recorded from the output of the retinal ganglion cell layer in a larval tiger salamander retina. For a full description of recording preparation see [52]. These data can be found at https://doi.org/10.5061/dryad.4qrfj6qm8.

### B. Visual stimuli: checkerboard noise and natural movies

A white noise checkerboard stimulus consisting of binary black or white squares, was presented at 30 frames per second (fps) for 30 minutes both before and after the natural scene presentations. Following this, five distinct 20-second natural movies were shown in a pseudorandom order, with each movie being displayed at least 80 times. Specifically, the tree, water, grasses, fish, and optic flow movies were repeated 83, 80, 84, 91, and 85 times, respectively. While all natural scenes were displayed at 60 fps, the tree stimulus was updated at 30 fps with each frame repeated twice to match the 60 fps rate. The optic flow movie was chosen for analysis because it elicited the strongest response.

### C. Low-Rank Ising model formulation and latent variable interpretation

Considering the energy for the joint distribution of visible binary spins *σ*_*i*_ and hidden Gaussian spins {*h*_*j*_}with mean 0 and unit variance. This yields a Hamiltonian for the joint

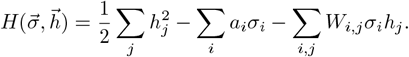

Marginalizing over hidden spins, gives the distribution over visible binary spins,

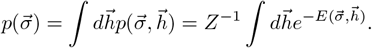

The distribution of visible spins after marginalizing out the hidden spins is also a Boltzmann distribution,

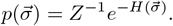

Equating these two expressions

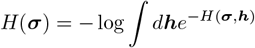

and expanding this out we obtain

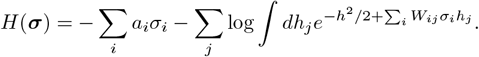

This expression can be greatly simplified with a cumulant generating function. Introducing zero mean unit variance Gaussians q_j_(h_j_) and the variable *t*_*j*_ = ∑_*i*_ W_i,j_σ_i_, which then gives

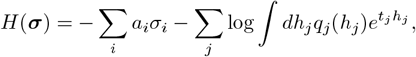

gives us the cumulant generating function

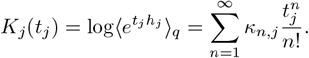

A mean 0 unit variance Gaussian, κ_1_ = 0, κ_2_ = 1 and all other terms are 0. Finally,

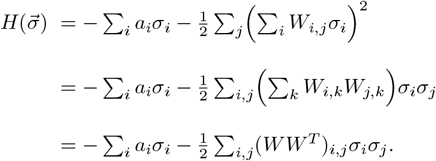

From this last expression it is clear that this is equivalent to an Ising model with interaction matrix *J*_*i,j*_ = (*W W* ^*T*^ )_*i,j*_, where the interaction matrix is rank k by construction.

### D. Inference of low-rank and full-rank Ising models

Ising model inference was done using factored minimum probability flow [86, 87]. As this method depends on model samples, sampling was doing using block Gibbs sampling in all cases, using 200 steps of sampling. This parallel update scheme is important for rapid mixing of Markov chains. Optimization was performed in PyTorch using LBFGS optimizer. An 80/20 train test split was used for cross validation. For the noise stimulus spiking activity was shuffled in one second chunks to discourage test set contamination. For the repeated natural scene data, spiking data was shuffled by trial index.

### E. Estimating the partition function for sparse retinal activity

Ising models in general have intractable partition functions. As retina activity is sparse the Good-Turing estimator is used [88] to infer missing mass which is then used to compute the missing portion of energy in the energy-based models. This method is explored in [89], and found to out perform other methods of partition function estimation for neural data. As an additional check upper and lower bounds on the log partition function are computed using a tractable model defined by independent neurons and the Bogoliubov bounds [90]. In all cases considered the upper and lower bounds are tight around the Good-Turing estimate. Together this suggests estimates of the partition function are accurate.

### F. Estimating receptive fields for neurons and latent variables

Latent variable receptive fields are estimated using linear regression on the noise stimulus. One second of the noise stimulus is concatenated to predict the activity of each latent variable at time t. Cross validation was used to determine the rank of the space-time filters. Again, the data was shuffled in one second chunks to avoid test set contamination. In all cases the rank-one fully space-time separable model was found to be optimal for each latent variable (Fig. S11). Performance on held out test data for all latent variables is shown in figure (Fig. S12). Performance declines as a function of latent variable index suggesting latent variables become noisier and more weakly coupled to the stimulus as a function of latent variable index. To estimate spatio-temporal filters for individual cells we use logistic regression where the weight matrix is constrained to be rank-one. Weights are initialized with the spike triggered average spatio-temporal filters and then optimization is done using LBFGS in PyTorch. Models are regularized using l2 regularization where the parameter was selected by cross-validation using the same approach as described for the linear regression spatio-temporal filters.

### G. Estimating nonlinear models

A more flexible generalized additive model with identity link is also considered. Here the filter inferred from linear regression is also allowed a spline nonlinearity using 20 B-spline basis functions. Smoothness was encouraged by a penalty on the second-derivative. At the limit of infinite regularization this will return the linear model. The smoothing parameter was selected using the same cross-validation procedure as the linear model. The inferred nonlinearities are given (Fig. S13), along with the performance on held out test data (Fig. S14). The leading latent variable shows the largest degree of nonlinearity, and the largest improvement from including the nonlinearity. All other latent variables show nonlinearities closer to the linear model and much smaller improvements in model performance.

### H. Ising model rotation

We have described how the interaction matrix of the Ising model is given by *J*_*i,j*_ = *W W* ^*T*^ . The latent variable model then generates equivalent models when an orthogonal matrix *Q* such that *QQ*^*T*^ = *I* is applied, because *W QQ*^*T*^ *W* ^*T*^ = *W W* ^*T*^ . In the main text *Q* is chosen to make the latent variables uncorrelated and ordered by variance (Fig. S10). This is done by projecting the spiking data onto the latent variables, computing their covariance, and finding the matrix of eigenvectors *U* of this covariance matrix. The rotation matrix is then *W U*.

An approach based on diagonalizing the interaction *J*_*i,j*_ = *W W* ^*T*^ was also considered. Here, *J*_*i,j*_ is computed and a constant is added to the diagonal to ensure that the matrix is positive definite. The matrix of eigenvectors *U* and matr of eigenvalues Λ are computed. We then set 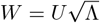

In our analysis of the variance maximizing choice of rotation, we find that the latent variables have stimulus sensitivity that declines as a function of latent variable index. To understand how similar the variance maximizing latent variables are to maximally stimulus responsive latent variables we do Canonical Correlation Analysis between the latent variable activity and the stimulus (CCA). This defines how to rotate hidden unit activity to be ordered by maximal correlation with the stimulus (Fig. S16). The last rotation considered is the CCA between the latent variable activity at time *t* and the latent variable activity at time *t* + 1. This identities features by how useful they are for predicting future stimulus states (Fig. S17).

In addition, there is a degree of freedom in the diagonal of the interaction matrix *J*_*ij*_. The diagonal of the matrix can be changed and any resulting modification to the energy function can be undone by modifying the bias term. As modifications to the diagonal are only guaranteed to preserved eigenvectors when a constant shift is added to each entry, this degree of freedom must be fixed before analyzing the matrix *W* . We select the diagonal that minimizes the nuclear norm of *J*_*ij*_.

### I. Natural image and natural scene masking and dataset curation

Both the CIFAR image datasets and the Action400 video datasets were converted to grayscale and resized to have the same resolution as the checkerboard stimulus (40 pixels by 40 pixels). For the Action400 videos, if the aspect ratio was not square the center crop was used. Images and videos were then scaled to be in the range [0, 1]. Finally, images and videos were masked to have activity in approximately the same region as the retina population response.

To mask images and videos the spatial receptive field of the first latent variable is taken and approximated with a two dimensional isotropic gaussian with parameters for spread, x-center, y-center, and amplitude. Parameters were estimated using the scipy implementation of the Levenberg-Marquardt algorithm. The mask was then normalized to have a maximum value of one. This mask was then manipulatively applied to the image. Noise was then added in the same way with the inverse mask, defined as 1 − mask. The noise distribution was uniform on [0, .05]

The CIFAR datasets were used without further processing or curation. The Action400 dataset only used videos with a framerate of 30 fps.

### J. Stimulus reconstruction and prediction

For the stimulus reconstruction task Action400 frames were projected onto the different bases and then reconstructed using linear regression from this lower dimensional subspace. l2 regularization was used, and the regularization strength was selected via cross validation. The same approach was used for the stimulus prediction task.

In the stimulus prediction task the optimal model was constructed using reduced rank regression. For a recent review focused on applications to neuroscience see [91]. For the multivariate regression task of predicting several different pixels, multiple different performance measures were considered. The standard definition of r^2^ was used for individual pixels 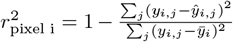A global measure of r^2^ was defined as 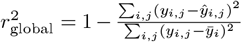. For the principal component measure of r^2^ both data and estimates were projected onto the natural image principal components and r^2^ was computed as 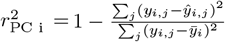

---

**FIG. S1.**
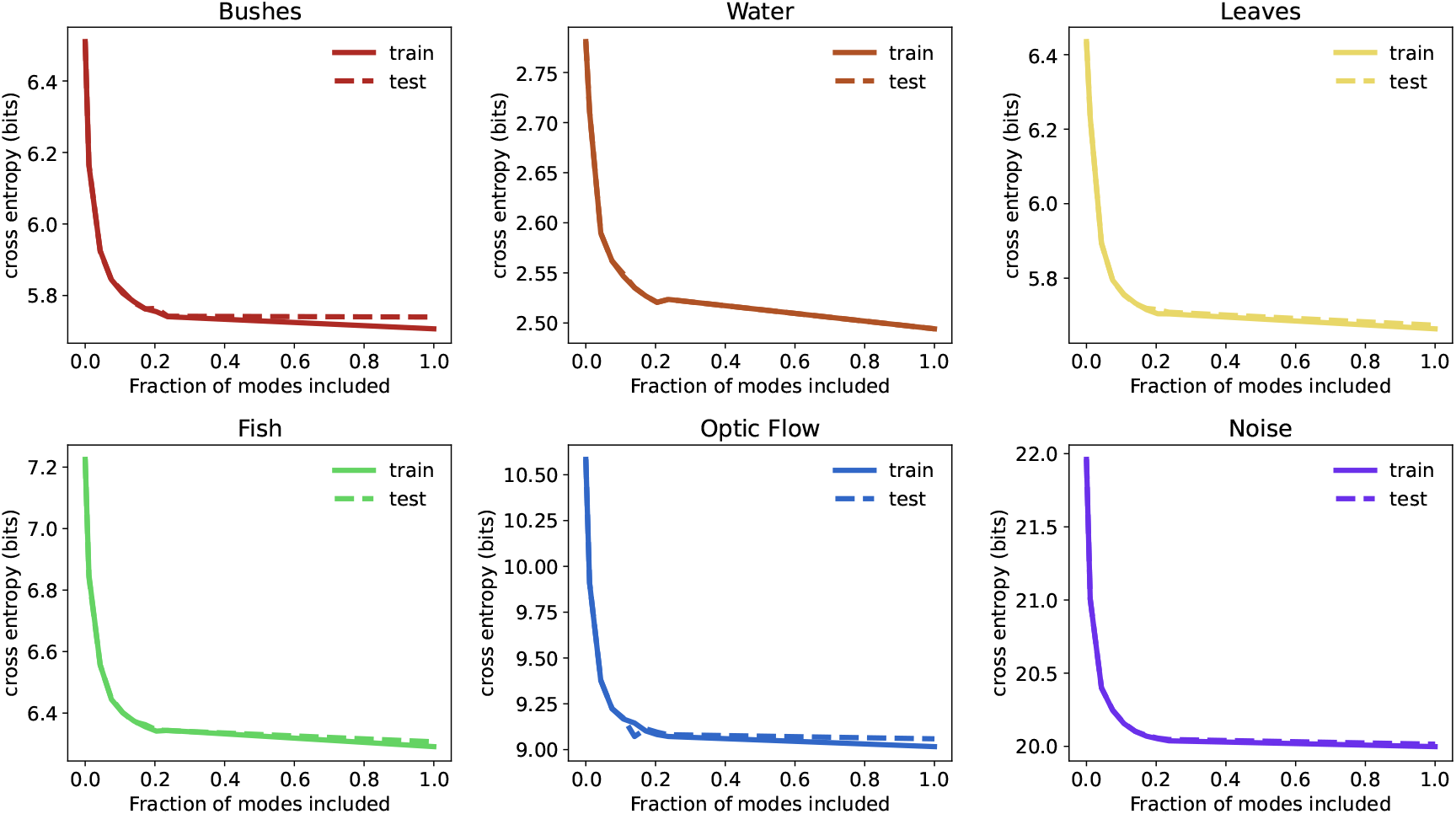
Cross entropy is shown as a function of rank for all videos. Performance quickly saturates across all videos, including naturalistic and artificial stimuli

**FIG. S2.**
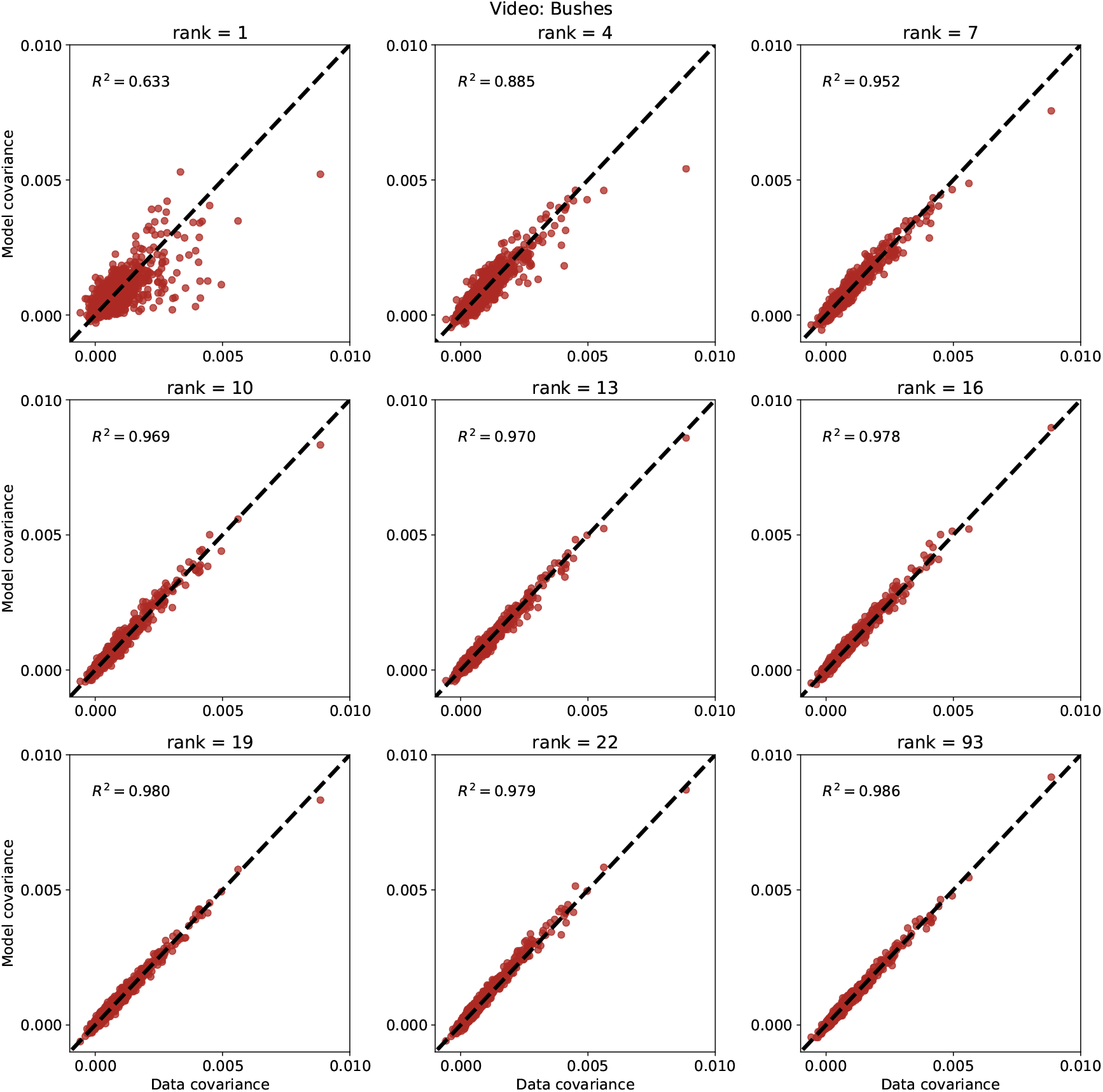
The relationship between model covariance and data covariance is shown for all cells for a range of numbers of latent variables. (Bushes video)

**FIG. S3.**
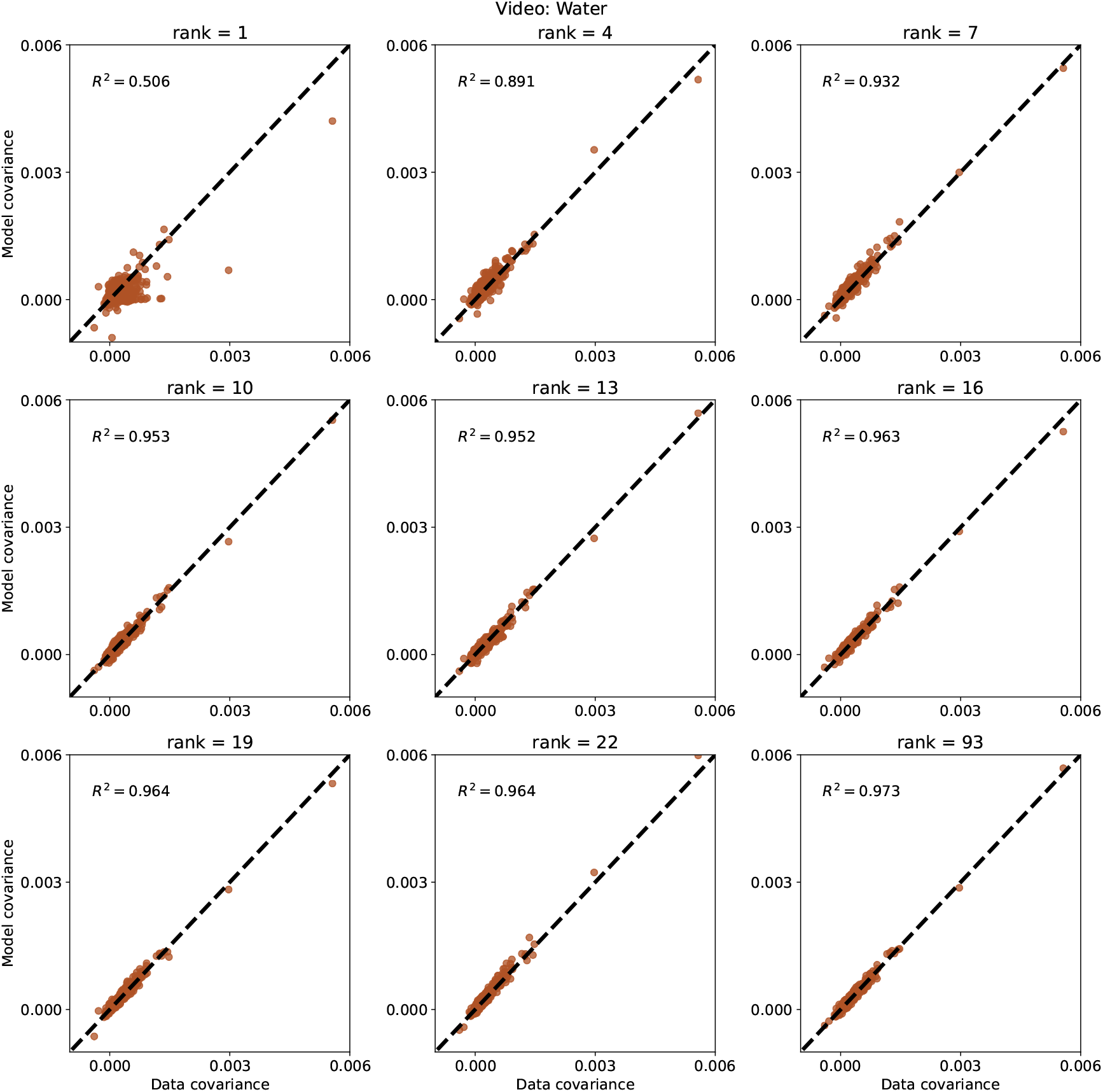
he relationship between model covariance and data covariance is shown for all cells for a range of numbers of latent variables. (Water video)

**FIG. S4.**
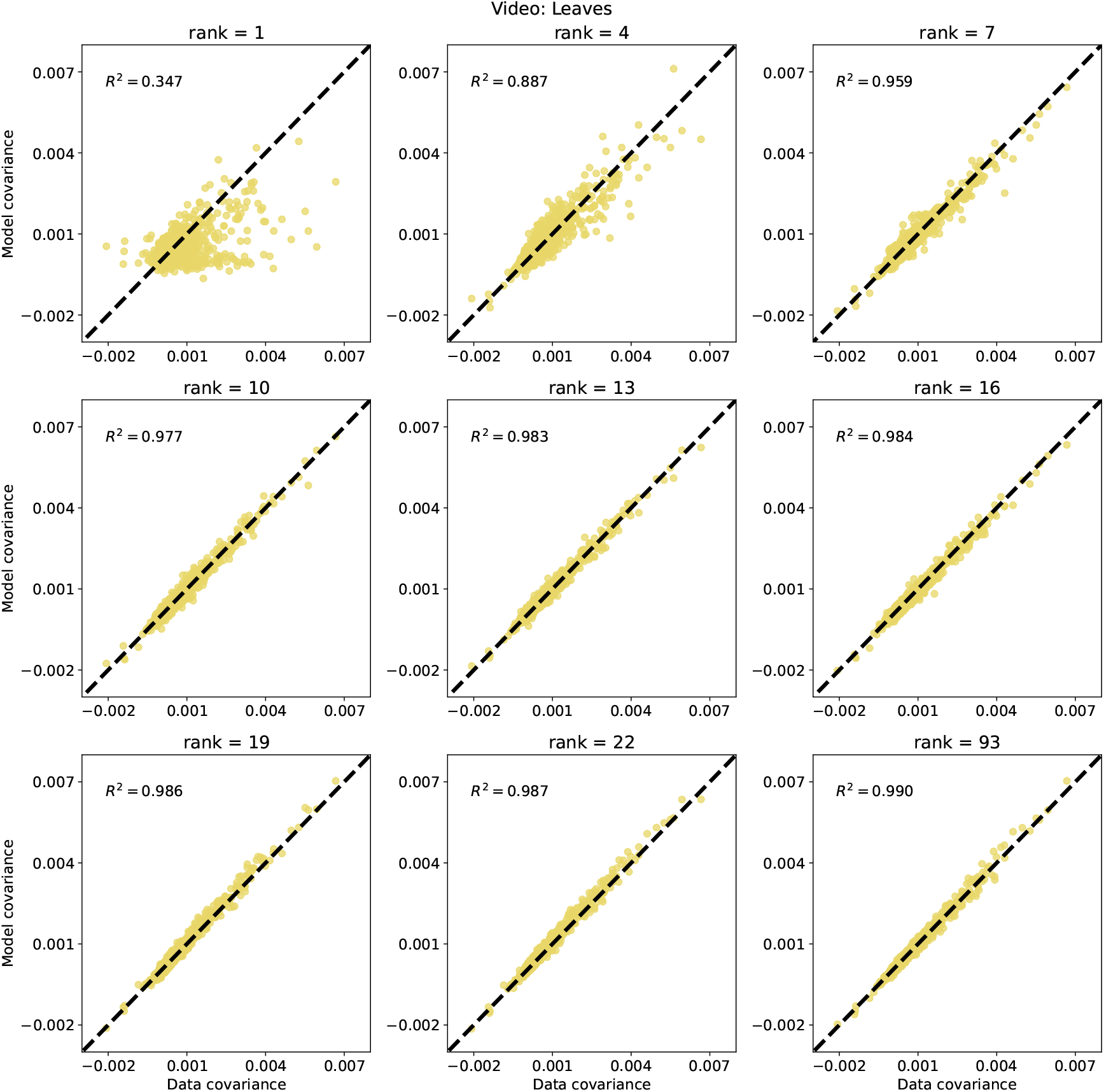
he relationship between model covariance and data covariance is shown for all cells for a range of numbers of latent variables. (Leaves video)

**FIG. S5.**
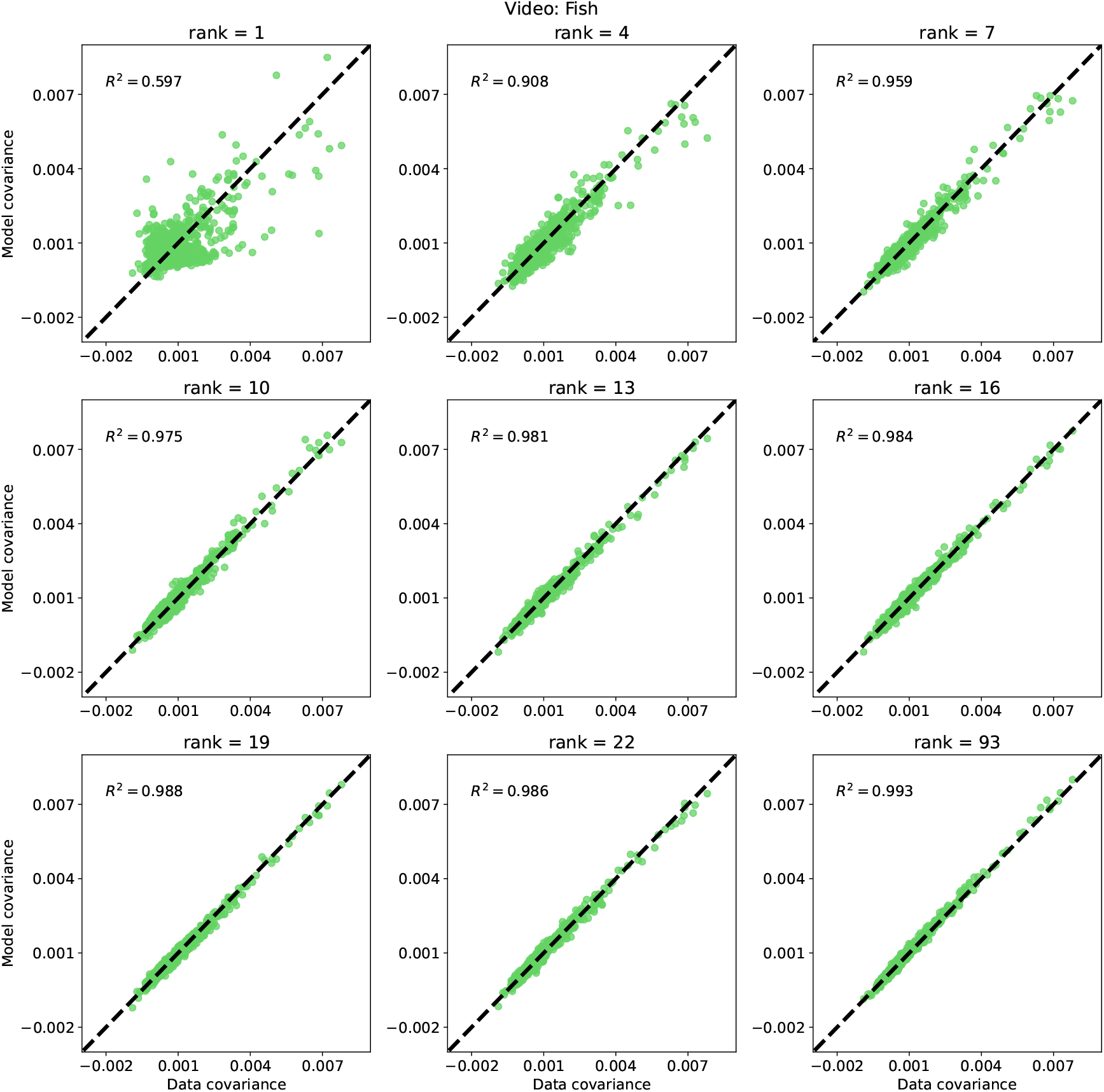
he relationship between model covariance and data covariance is shown for all cells for a range of numbers of latent variables. (Fish video)

**FIG. S6.**
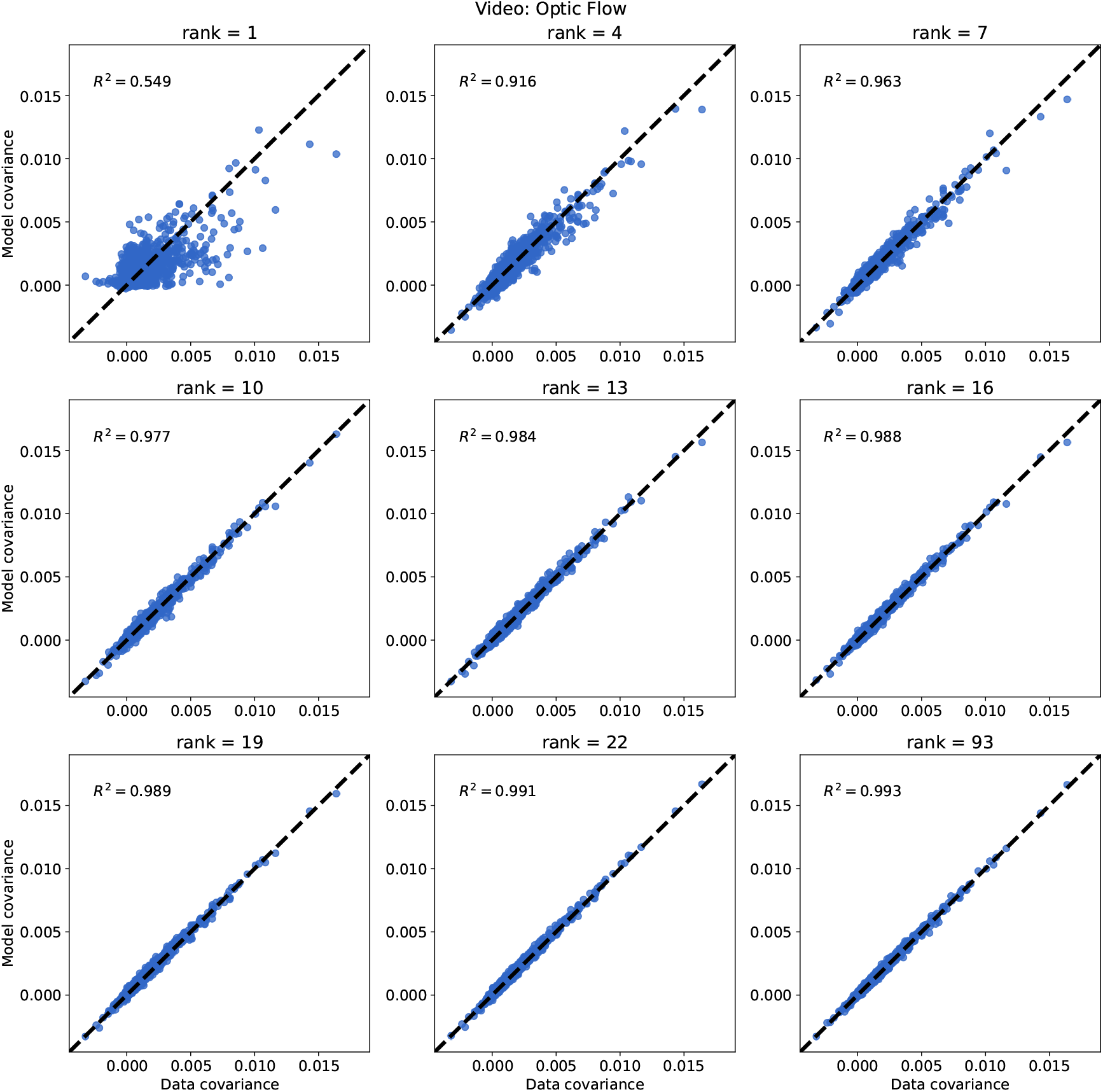
he relationship between model covariance and data covariance is shown for all cells for a range of numbers of latent variables. (Optic Flow video)

**FIG. S7.**
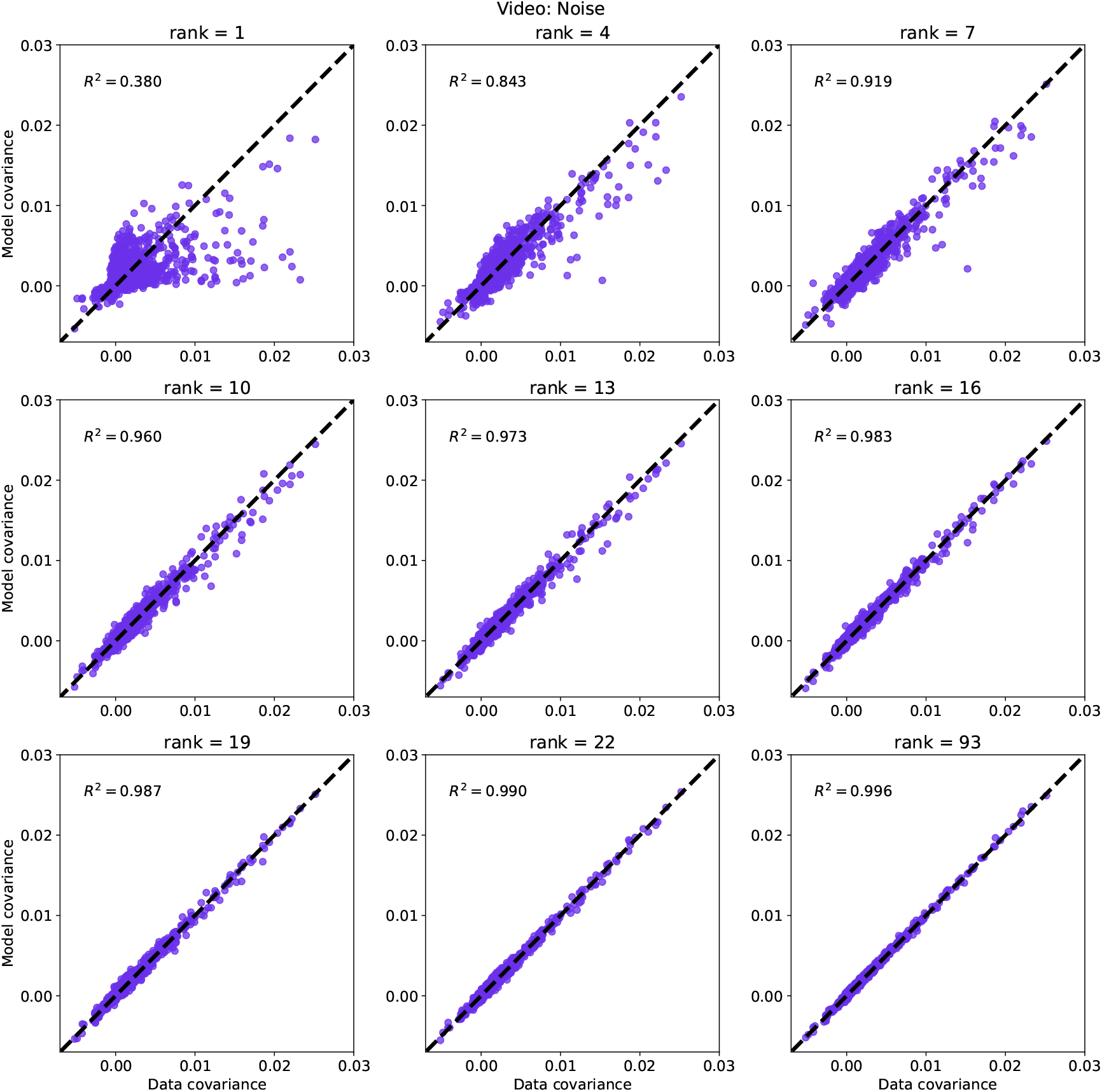
he relationship between model covariance and data covariance is shown for all cells for a range of numbers of latent variables. (Noise video)

**FIG. S8.**
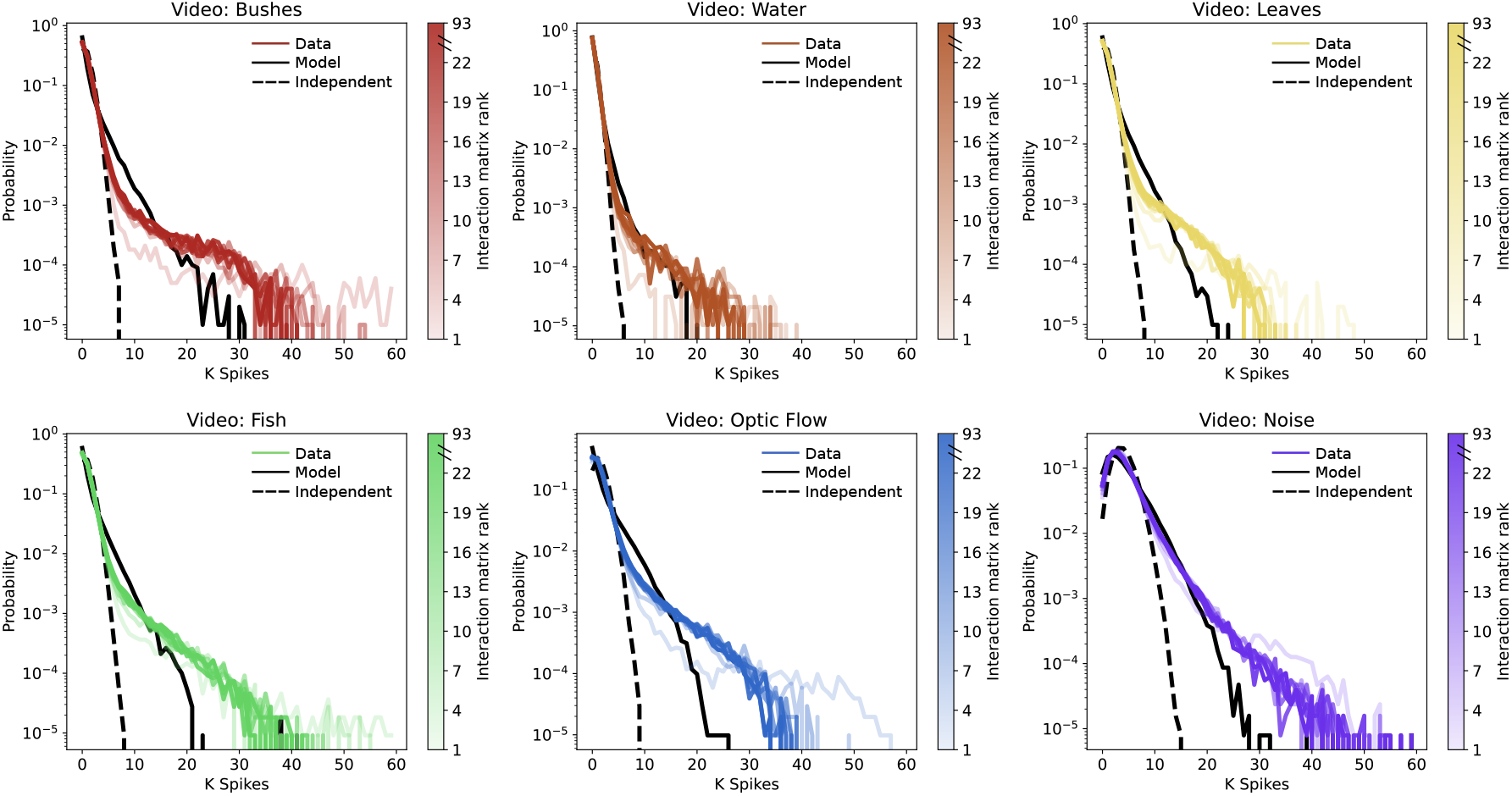
Distribution of spike counts in a time bin for data, Ising models with variable rank, and the rate-matched independent model are shown for all stimulus conditions.

**FIG. S9.**
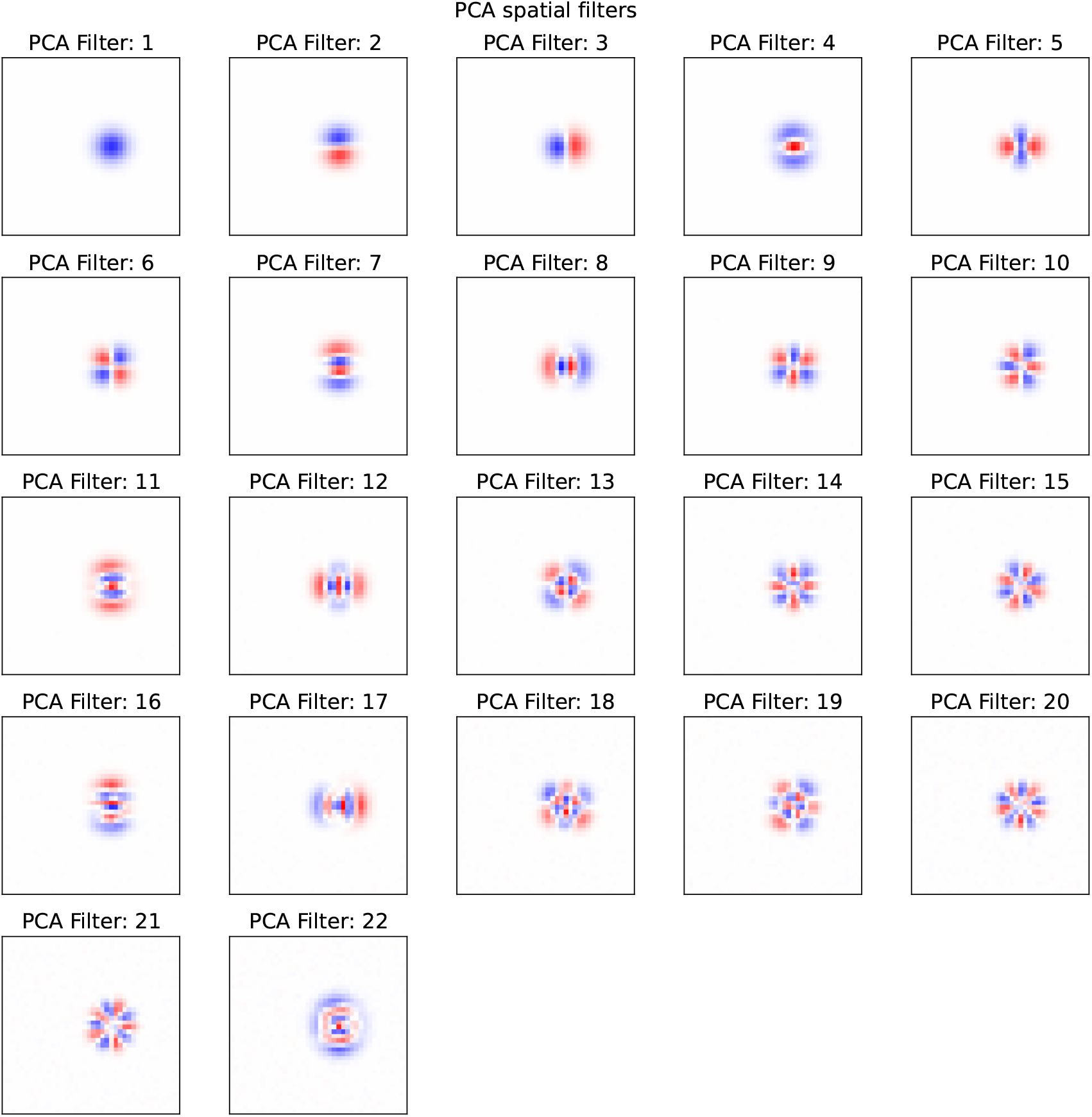
The first 22 principal components for the masked natural images dataset are shown.

**FIG. S10.**
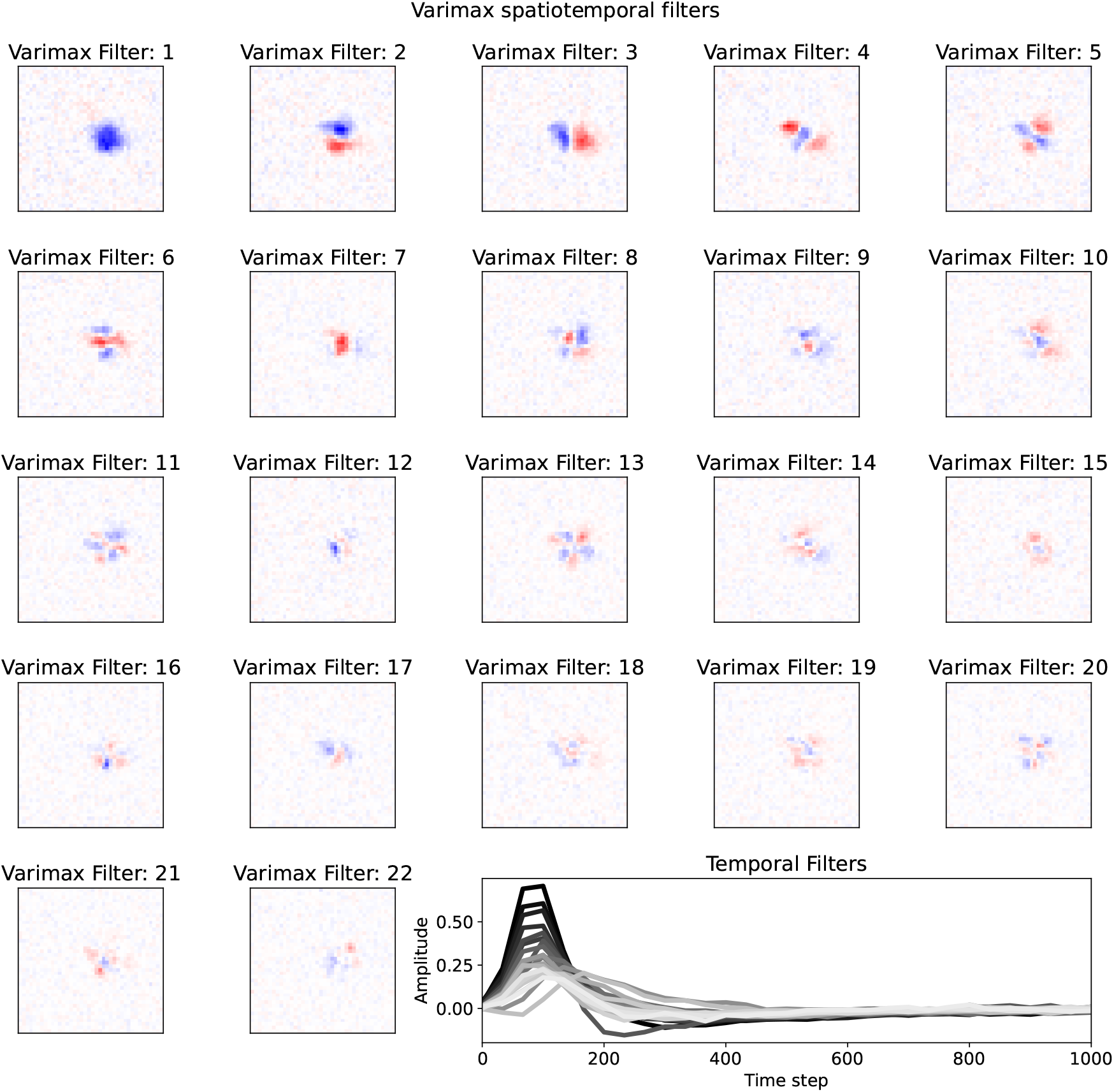
The first 22 retina spatial and temporal filters are shown for the variance maximizing rotation matrix. Temporal filter index decreases from dark to light

**FIG. S11.**
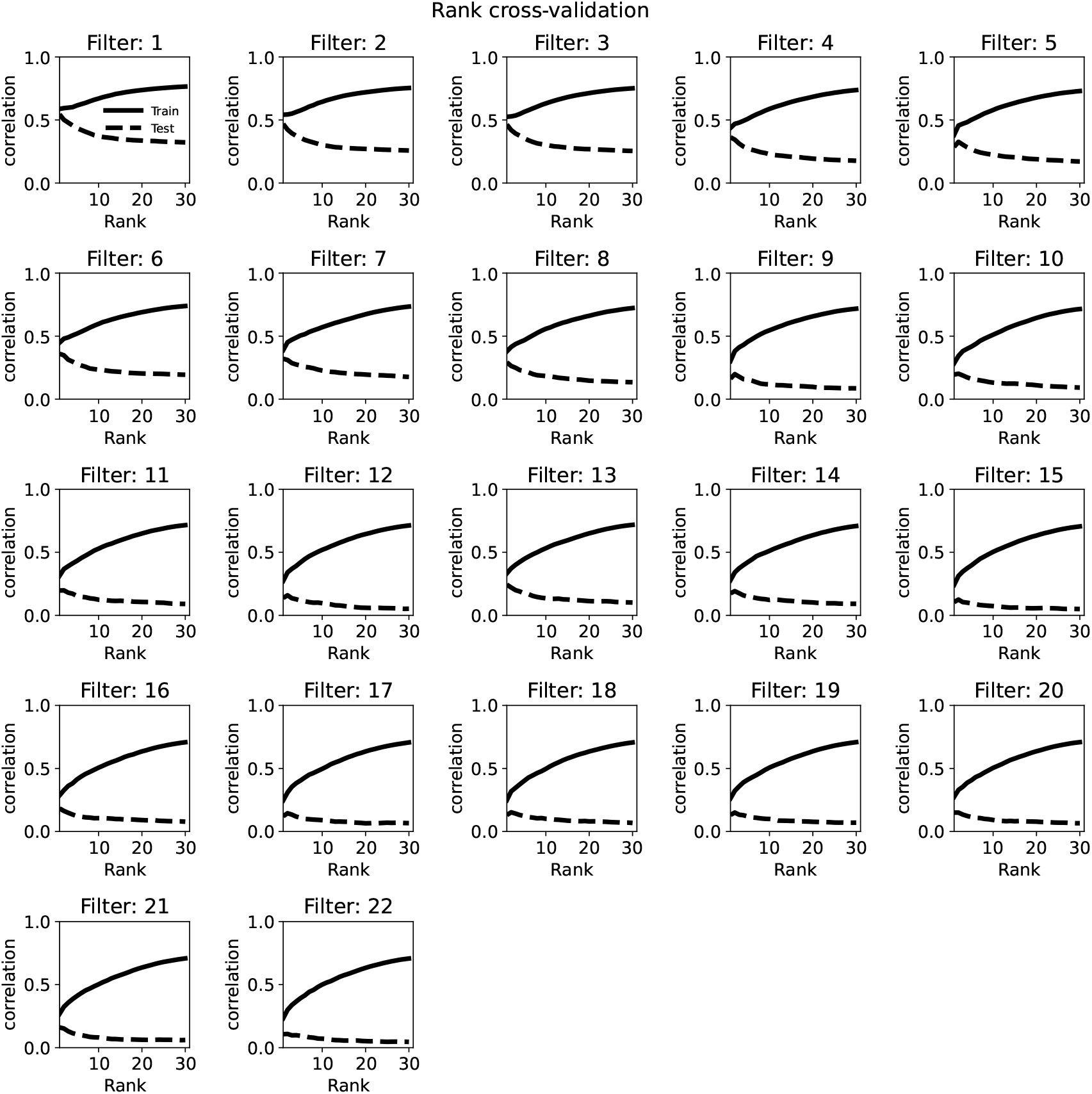
Cross validation performance for all 22 retina filters as a function of rank for decomposition of spatiotemporal coefficients. Rank 1 corresponds to fully space-time separable filters

**FIG. S12.**
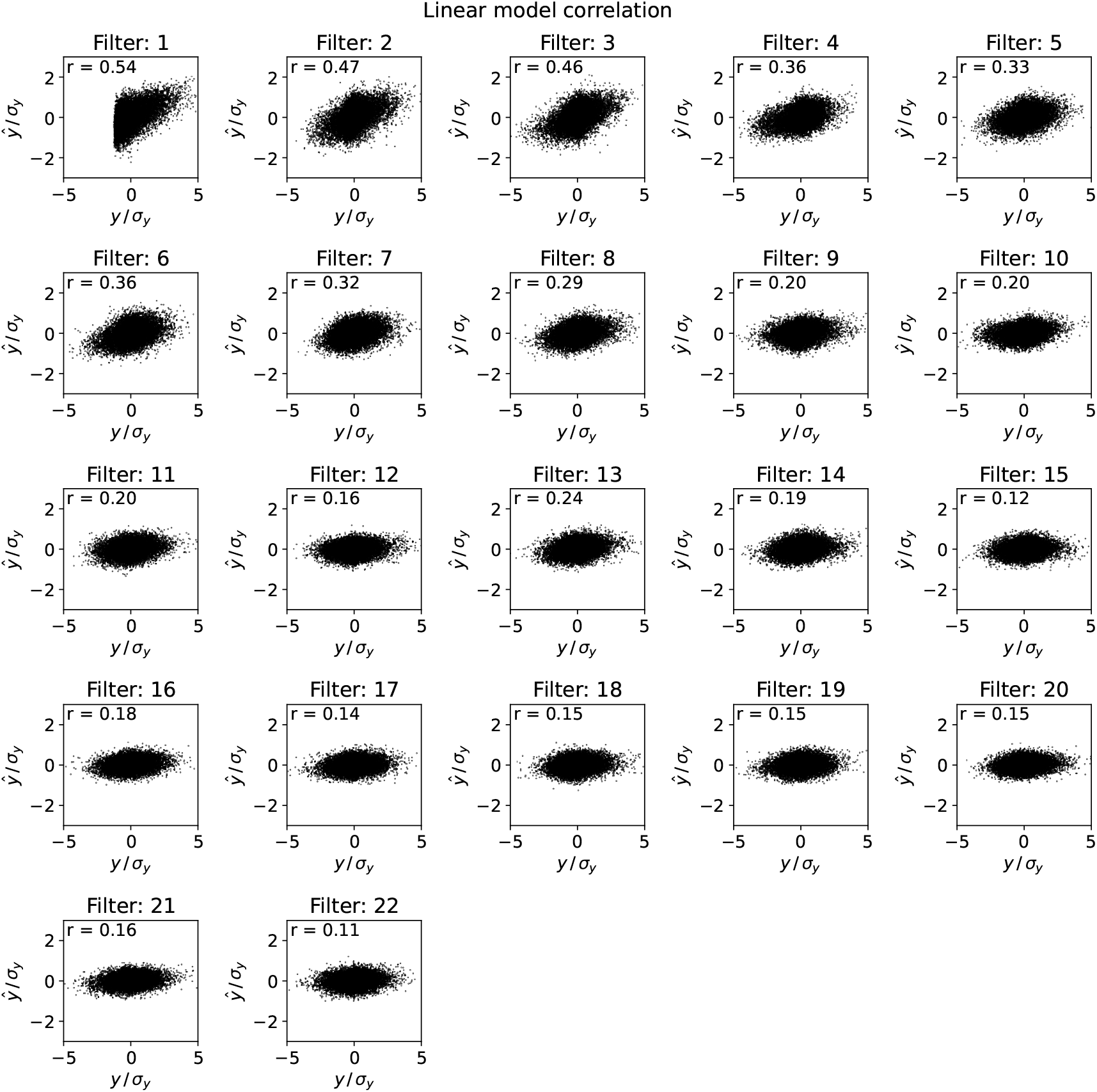
Scatter plots showing correlation between true values of latent variable activity for held out test data and model estimates for all latent variables.

**FIG. S13.**
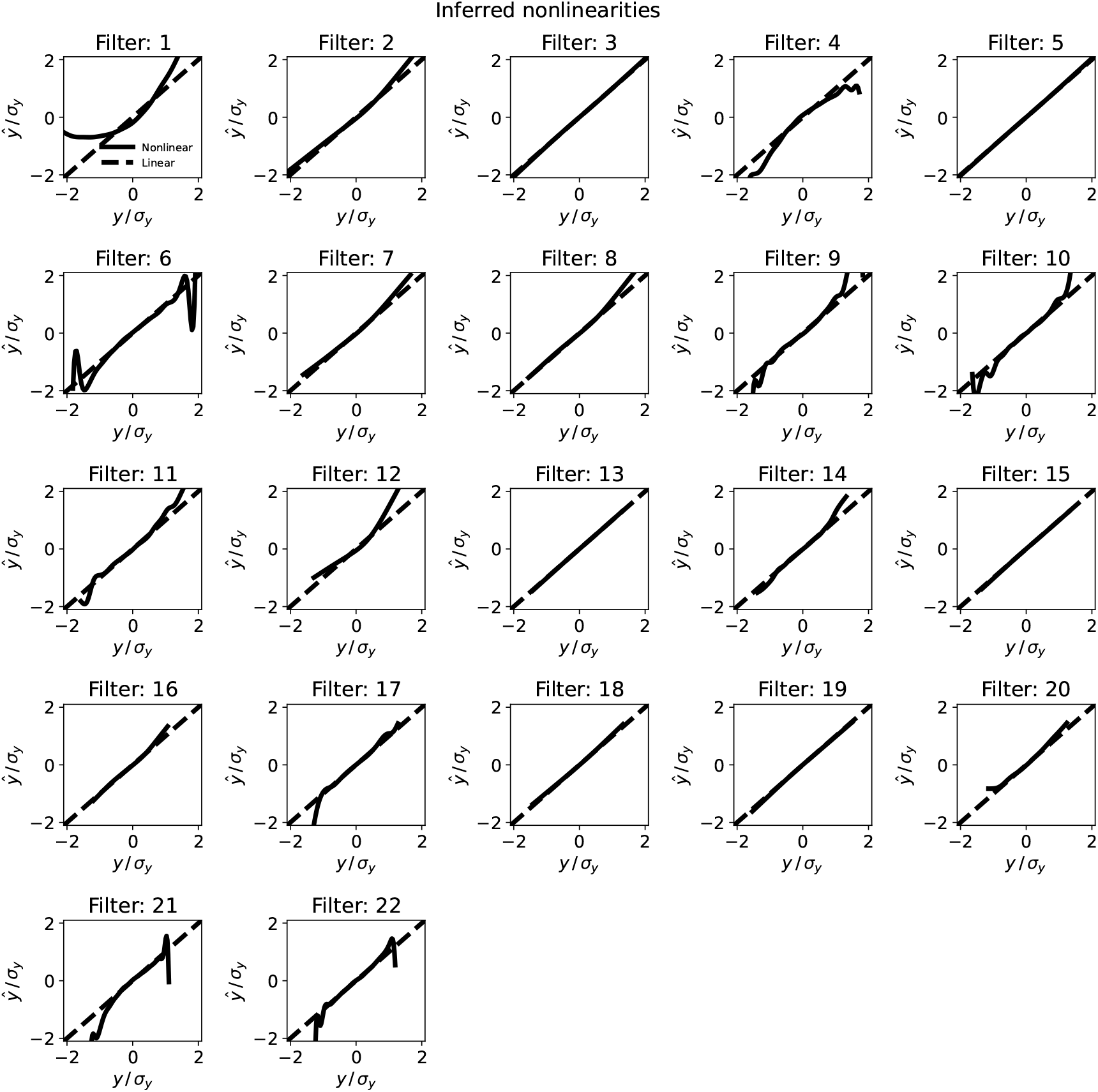
Inferred nonlinearities for each latent variable encoding model. The leading filter shows the strongest dependence on an added nonlinearity.

**FIG. S14.**
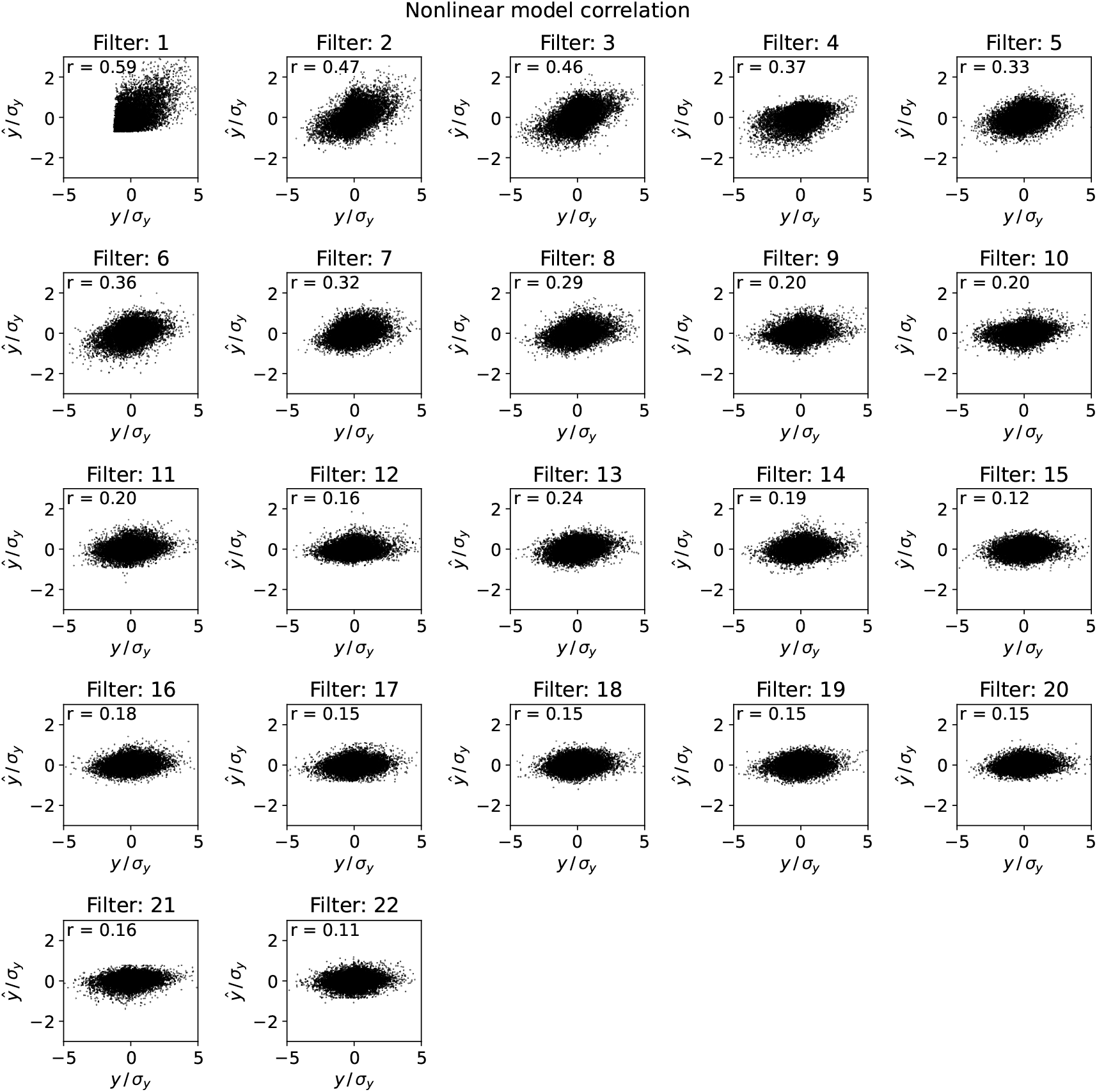
Scatter plots showing correlation between true values of latent variable activity for held out test data and model estimates for all latent variables with the addition of a nonlinearity. Only the filter 1 shows significant improvement.

**FIG. S15.**
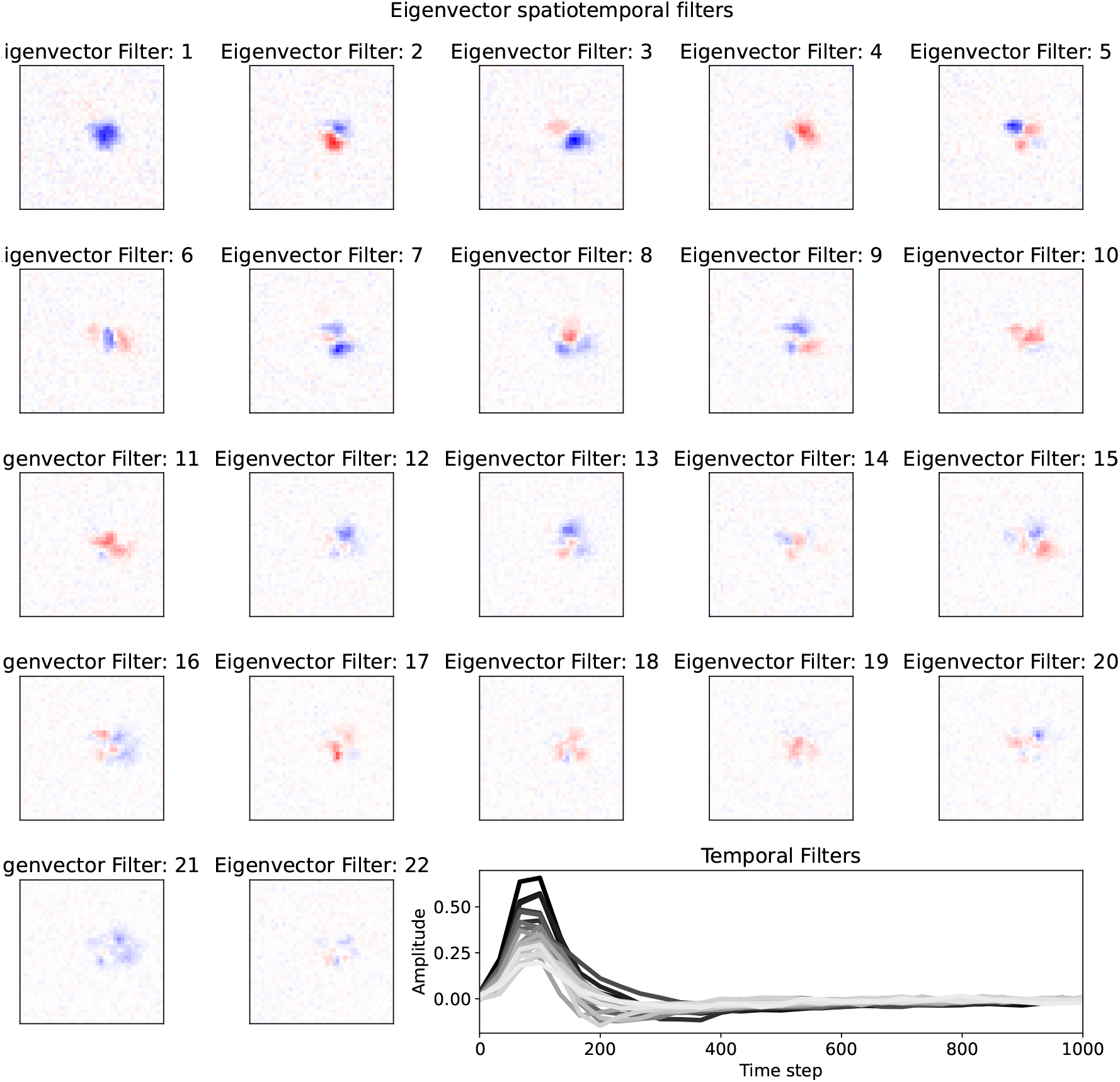
Multiple possible choices of rotation matrix *U* are considered for the inferred Ising models and lead to similar inferred filters. Spatial and temporal filters are shown for all filters where the rotation matrix was chosen to diagonalize the interaction matrix *J*_*i,j*_

**FIG. S16.**
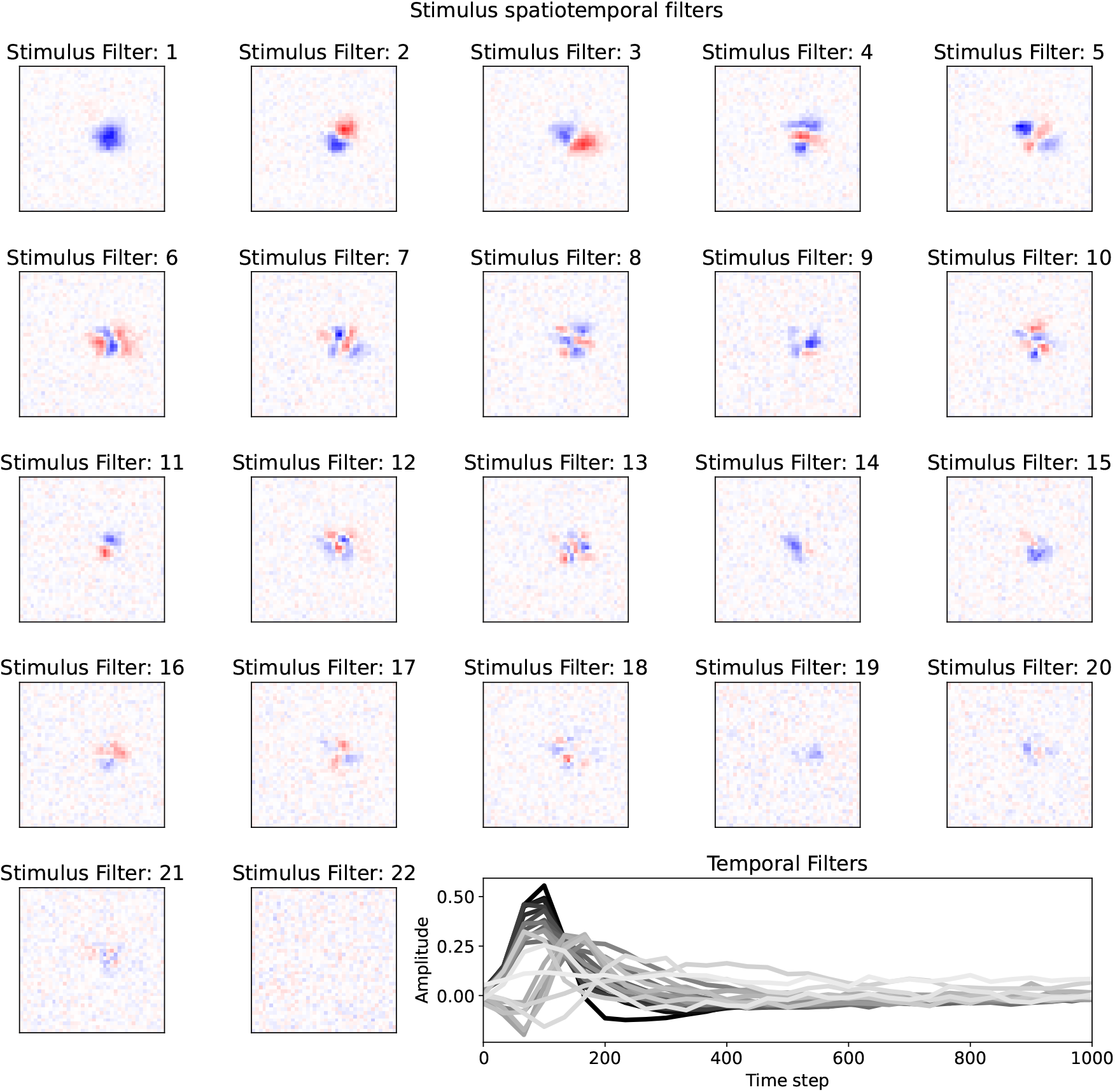
Spatial and temporal filters are shown for all filters where the rotation matrix was chosen to diagonalize the cross-covariance between latent variable activity and the stimulus. This orders latent variables by how strongly they are coupled to the stimulus.

**FIG. S17.**
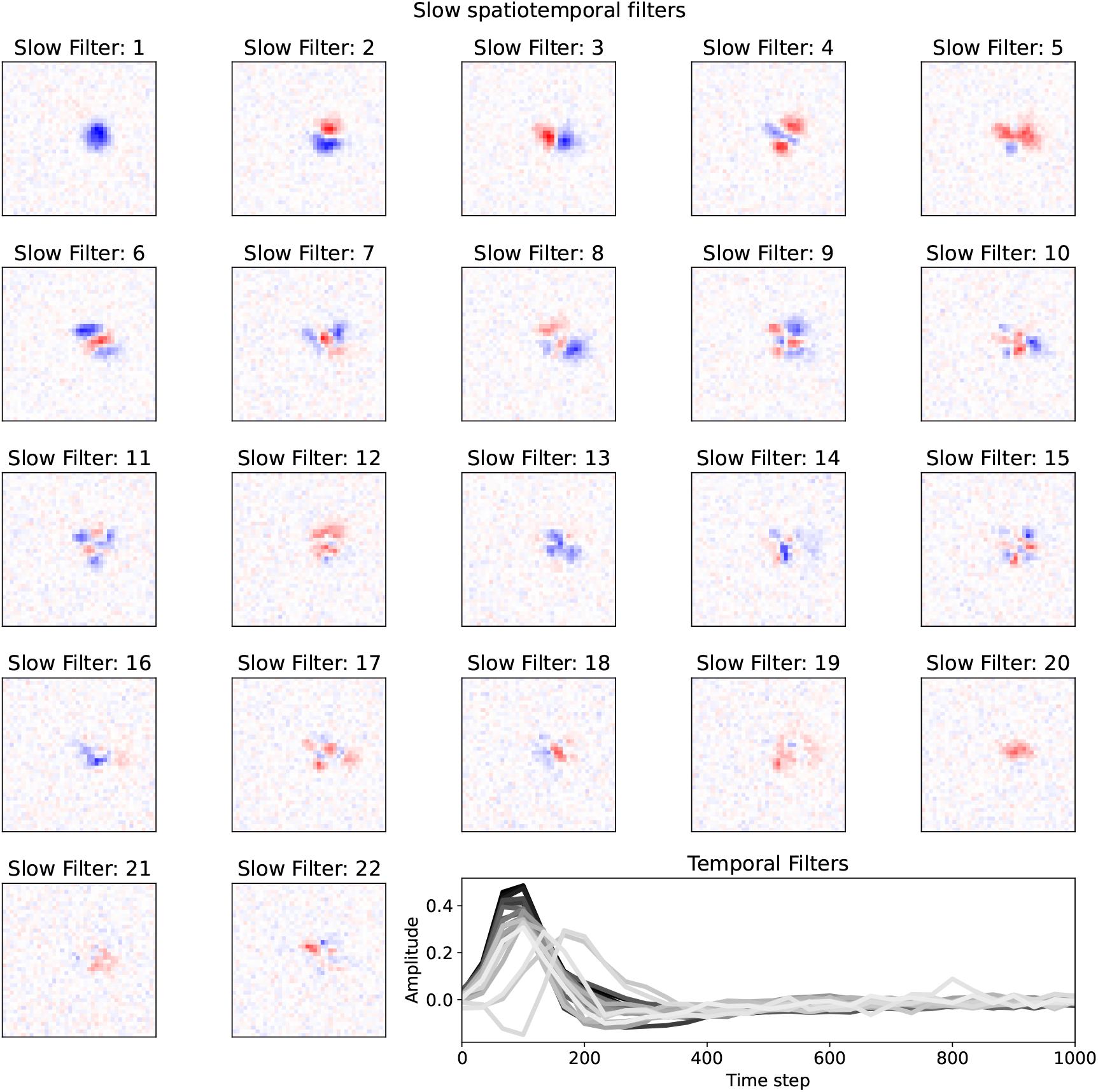
Spatial and temporal filters are shown for all filters where the rotation matrix was chosen to diagonalize the cross-correlation between latent variable activity at time *t* and time *t* + 1. This orders latent variables by how useful they are for predicting the next step in time. The top several filters are similar across many different choices of rotation matrix.

**FIG. S18.**
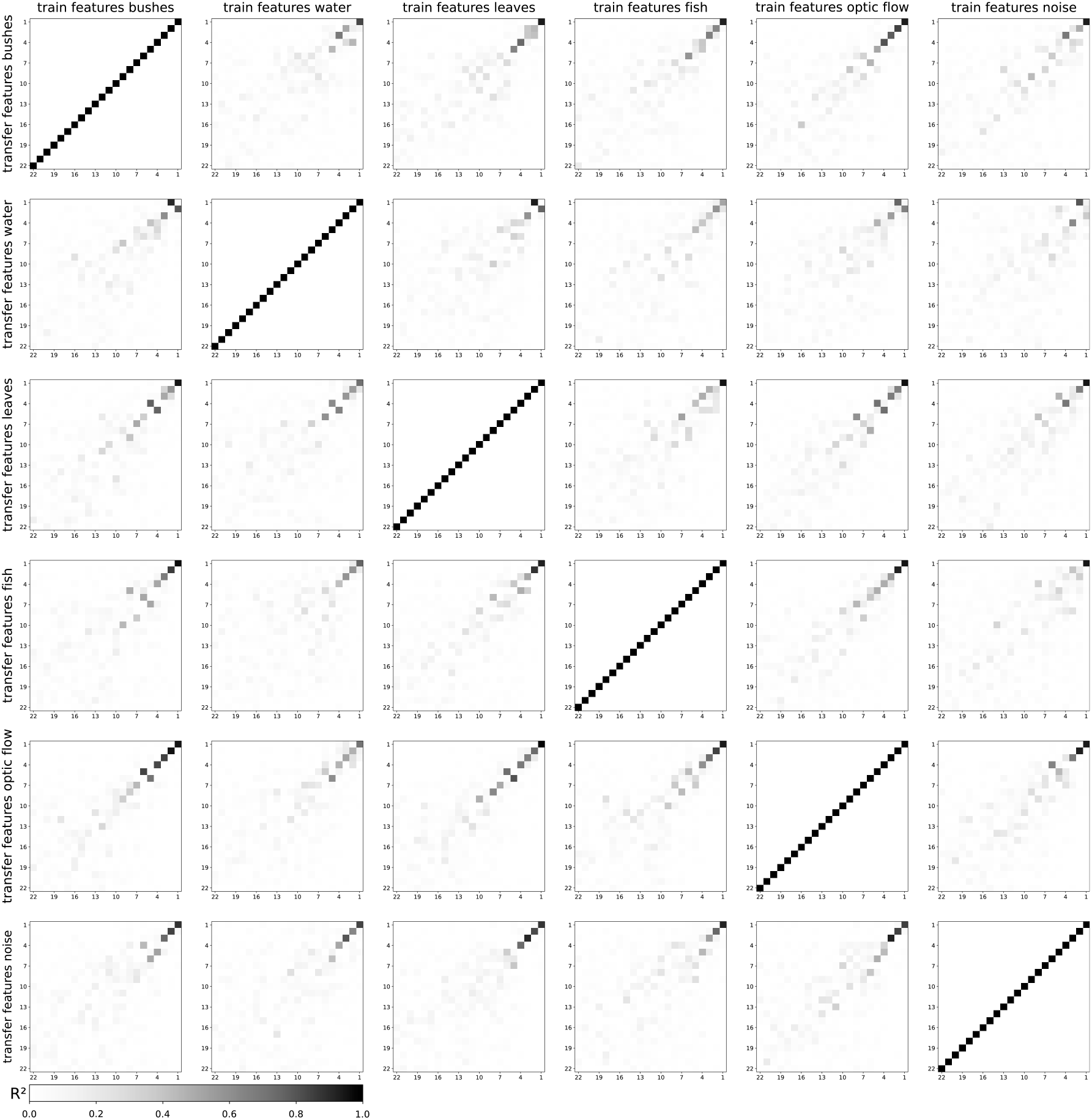
Stability of top three latent variables is consistent across stimulus conditions. The transfer learning task performance is shown for all pairwise comparisons.

**FIG. S19.**
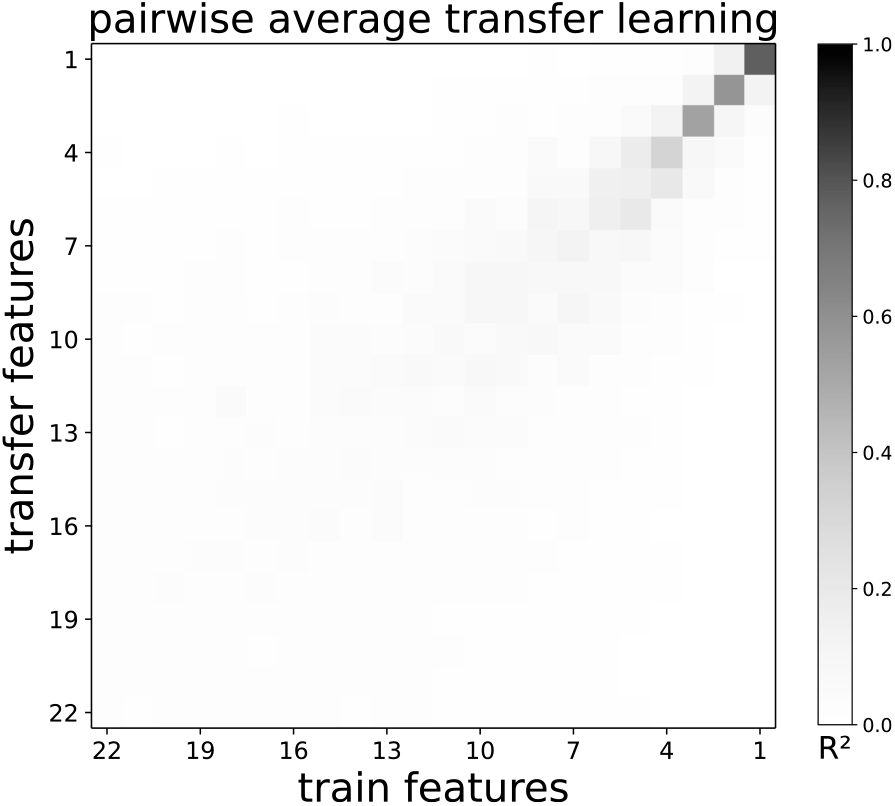
Average performance over all pairwise transfer learning performances show the top three latent variables are particularly stable. Similarity decays as a function of latent variable index.

**FIG. S20.**
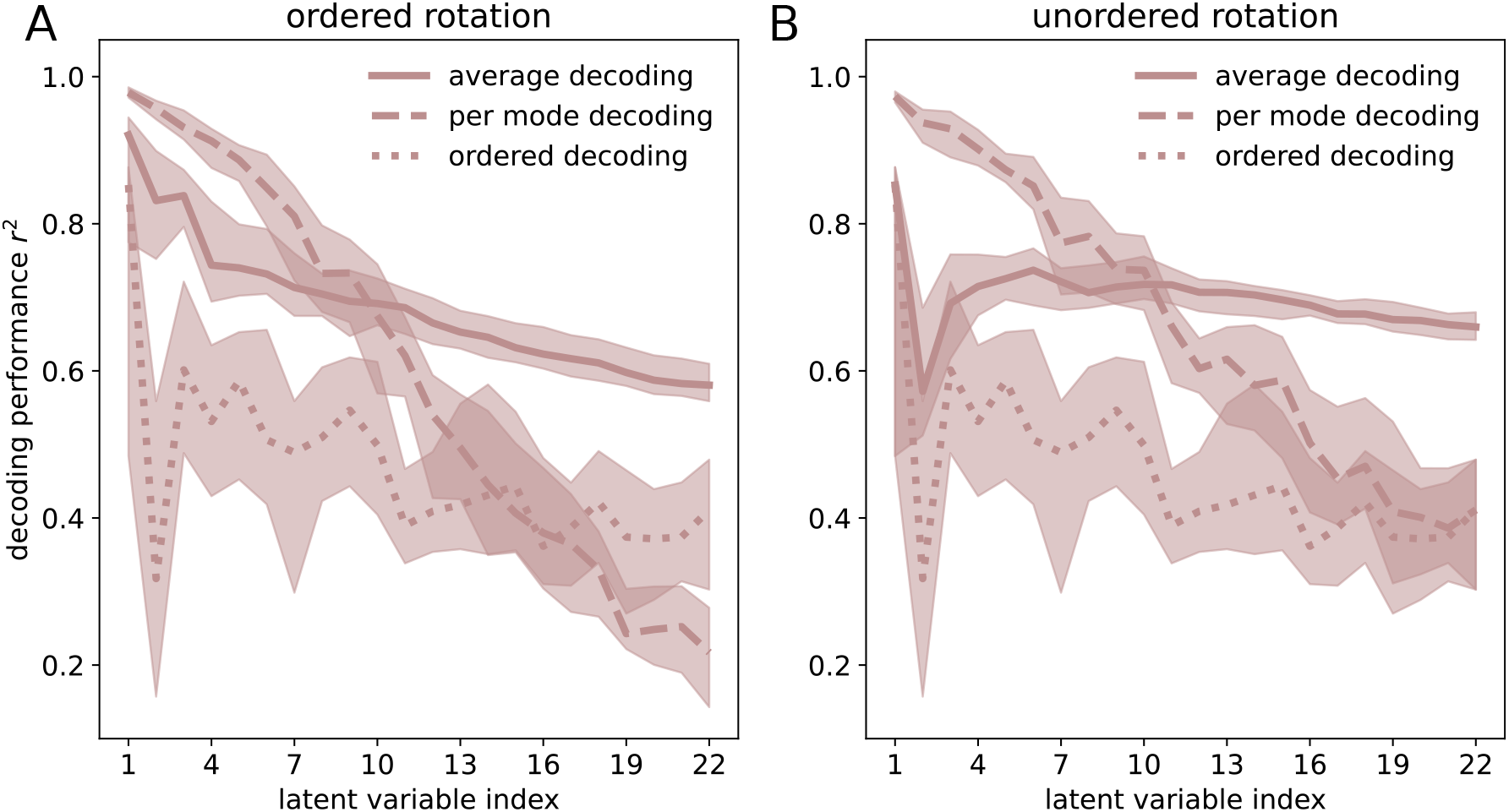
Decoding performance of average decoder, per mode decoder and ordered decoder are shown as a function of index for the transfer learning task. A) Ordered rotation chooses a rotation of the transfer weight matrix that maximizes the variability under the training spikes. B) Unordered rotation chooses a rotation of the transfer weight matrix that maximizes the variability under the transfer spikes. Both show decaying *r*^2^ as a function of mode index.

**FIG. S21.**
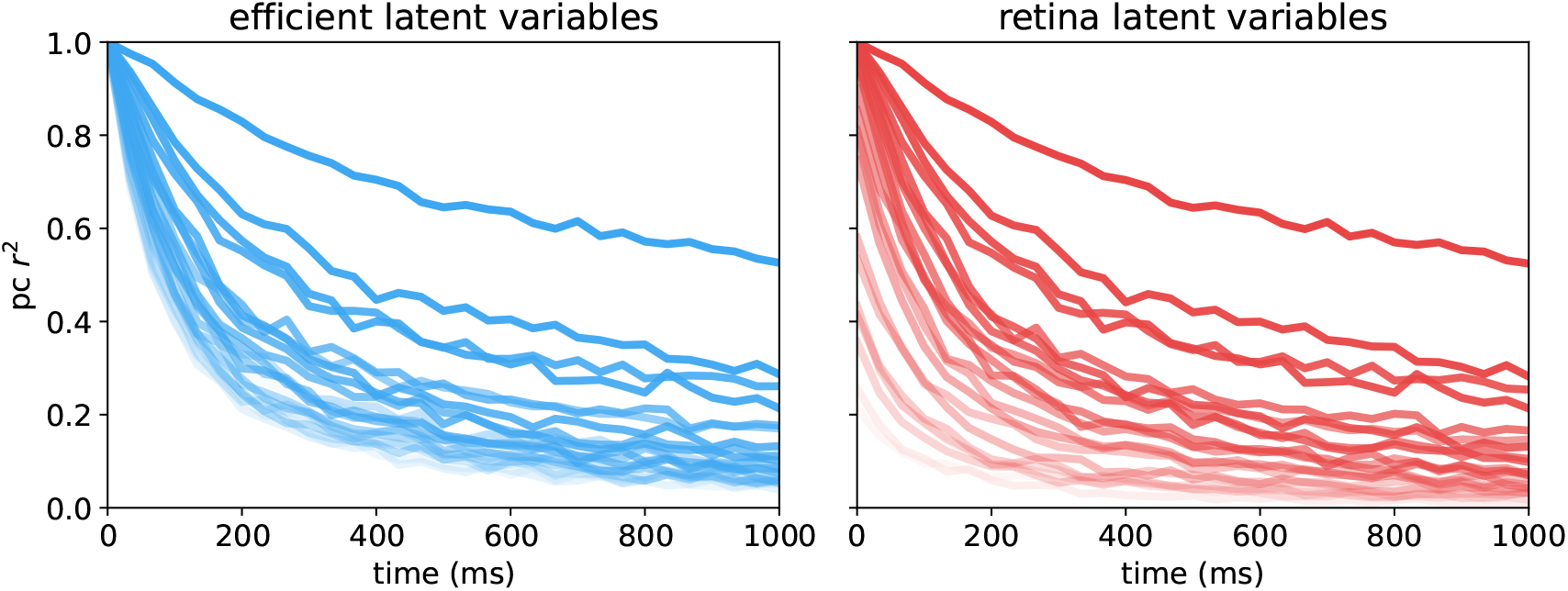
Latent-variable prediction dynamics for efficient and retinal codes. Prediction performance (*r*^2^) with natural image principal components is shown as a function of time for the efficient coding basis (left) and the retinal coding basis (right). Each curve corresponds to one principal component, ordered by decreasing variance, with opacity decreasing from the first to the twenty-second component. For both sets of latent variables, leading components retain predictive power over longer time scales, while higher-order components decay more rapidly.

**FIG. S22.**
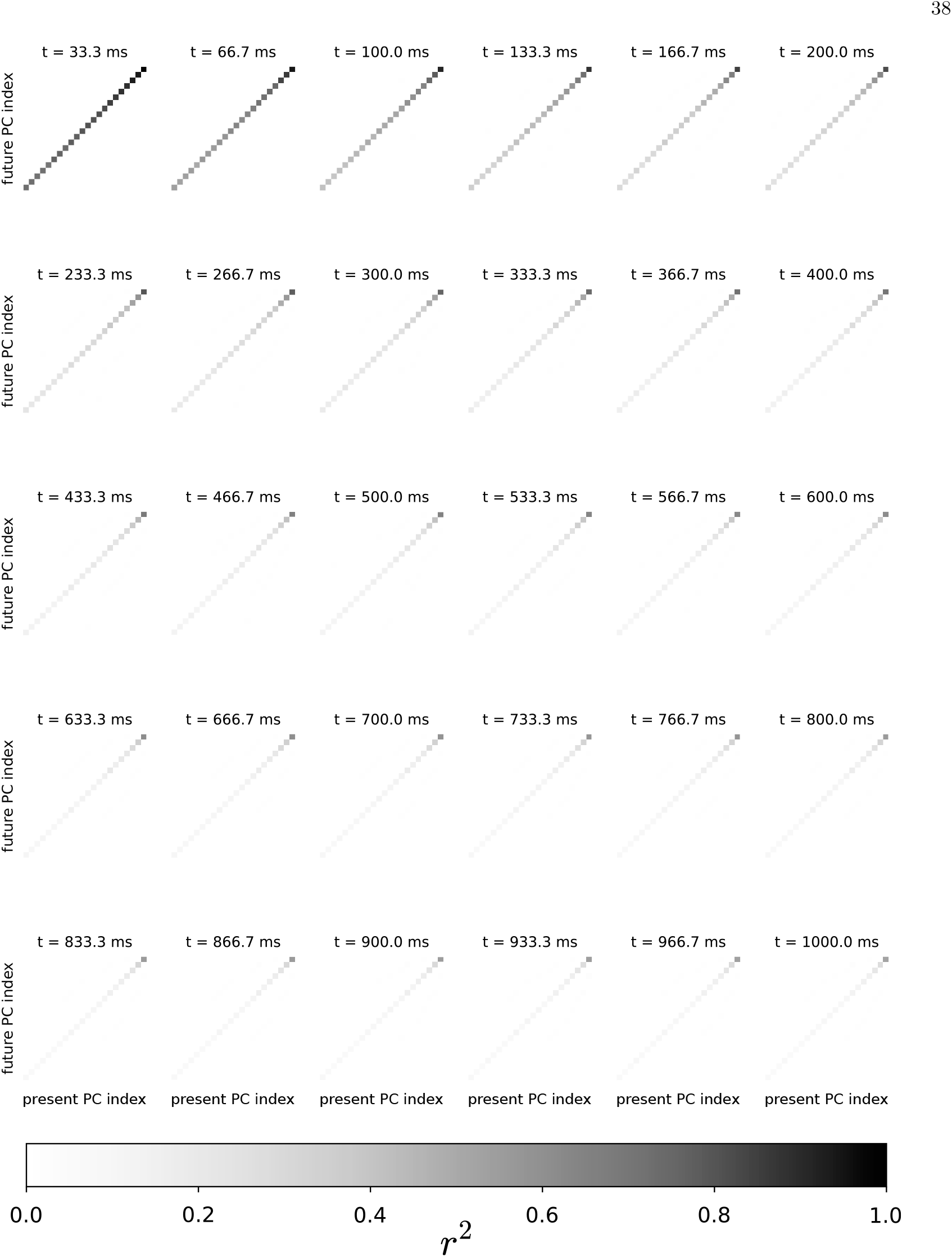
Natural scene principal components approximately diagonalize future states. Natural scenes are projected onto the first twenty-two principal components defined by the equal time covariance matrix. Across a range of times the squared correlation is reported between principal component *i* at time *t* and principal component *j* at time *t* + *k*. All off diagonal entries remain near zero across time.

## REFERENCE

[1] F. Attneave, Psychol. Rev. 61, 183 (1954), 13167245.

[2] H. B. Barlow, OUP Academic (2012), 10.7551/mitpress/9780262518420.003.0013.

[3] J. J. Atick and A. N. Redlich, Neural Comput. 2, 308 (1990).

[4] J. J. Atick and A. N. Redlich, Neural Comput. 4, 196 (1992).

[5] E. Doi, J. L. Gauthier, G. D. Field, J. Shlens, A. Sher, M. Greschner, T. A. Machado, L. H. Jepson, K. Mathieson, D. E. Gunning, A. M. Litke, L. Paninski, E. J. Chichilnisky, and E. P. Simoncelli, J. Neurosci. 32, 16256 (2012), 23152609.

[6] Y. Karklin and E. P. Simoncelli, Adv. Neural Inf. Process. Syst. 24:999–1007. (2011), 26273180.

[7] S. Roy, N. Y. Jun, E. L. Davis, J. Pearson, and G. D. Field, Nature 592, 409 (2021).

[8] N. Y. Jun, G. Field, and J. Pearson, Advances in Neural Information Processing Systems 35, 32311 (2022).

[9] X. Pitkow and M. Meister, Nat. Neurosci. 15, 628 (2012).

[10] V. Balasubramanian and M. J. B. I. I., Network 13, 531 (2002).

[11] S. Ocko, J. Lindsey, S. Ganguli, and S. Deny, Advances in Neural Information Processing Systems 31 (2018).

[12] H. Attias and C. Schreiner, Advances in Neural Information Processing Systems 9 (1996).

[13] M. S. Lewicki, Nat. Neurosci. 5, 356 (2002).

[14] E. C. Smith and M. S. Lewicki, Nature 439, 978 (2006).

[15] C. F. Stevens, Proc. Natl. Acad. Sci. U.S.A. 112, 9460 (2015).

[16] D. Zwicker, A. Murugan, and M. P. Brenner, Proc. Natl. Acad. Sci. U.S.A. 113, 5570 (2016).

[17] S. E. Palmer, O. Marre, M. J. Berry, and W. Bialek, Proc. Natl. Acad. Sci. U.S.A. 112, 6908 (2015).

[18] M. Chalk, O. Marre, and G. Tkacik, Advances in Neural Information Processing Systems 29 (2016).

[19] M. Chalk, O. Marre, and G. Tkačik, Proc. Natl. Acad. Sci. U.S.A. 115, 186 (2018).

[20] B. Liu, A. Hong, F. Rieke, and M. B. Manookin, Nat. Neurosci. 24, 1280 (2021).

[21] K. Bojanek, B. Lefebvre, J. M. Salisbury, O. Marre, and S. Palmer, bioRxiv , 2025.09.19.677348 (2025), 2025.09.19.677348.

[22] L. Taylor, F. Zenke, A. J. King, and N. S. Harper, bioRxiv , 2024.03.26.586771 (2024), 2024.03.26.586771.

[23] S. Klavinskis-Whiting, E. Fristed, Y. Singer, M. F. Iacaruso, A. J. King, and N. S. Harper, Curr. Biol. 35, 530 (2025).

[24] Y. Singer, Y. Teramoto, B. D. B. Willmore, J. W. H. Schnupp, A. J. King, and N. S. Harper, eLife (2018), 10.7554/eLife.31557.

[25] Y. Singer, L. Taylor, B. D. B. Willmore, A. J. King, and N. S. Harper, eLife (2023), 10.7554/eLife.52599.

[26] R. W. Williams, R. C. Strom, D. S. Rice, and D. Goldowitz, J. Neurosci. 16, 7193 (1996), 8929428.

[27] N. P. Shah, N. Brackbill, R. Samarakoon, C. Rhoades, A. Kling, A. Sher, A. Litke, Y. Singer, J. Shlens, and E. J. Chichilnisky, Neuron 110, 698 (2022), 34932942.

[28] P. W. Keeley, S. J. Eglen, and B. E. Reese, J. Comp. Neurol. 528, 2135 (2020).

[29] X. Ibarra-Soria, T. S. Nakahara, J. Lilue, Y. Jiang, C. Trimmer, M. A. A. Souza, P. H. M. Netto, K. Ikegami, N. R. Murphy, M. Kusma, A. Kirton, L. R. Saraiva, T. M. Keane, H. Matsunami, J. Mainland, F. Papes, and D. W. Logan, eLife (2017), 10.7554/eLife.21476.

[30] K. Rihani and S. Sachse, Front. Behav. Neurosci. 16:835680. (2022), 10.3389/fnbeh.2022.835680, 35548690.

[31] M. Bauer, W. Bialek, C. Goddard, C. M. Holmes, K. Krishnamurthy, S. E. Palmer, R. Pang, D. J. Schwab, and L. Susman, arXiv (2025), 10.48550/arXiv.2505.23398, 2505.23398.

[32] K. S. Brown and J. P. Sethna, Phys. Rev. E 68, 021904 (2003).

[33] R. N. Gutenkunst, J. J. Waterfall, F. P. Casey, K. S. Brown, C. R. Myers, and J. P. Sethna, PLoS Comput. Biol. 3, e189 (2007).

[34] J. J. Waterfall, F. P. Casey, R. N. Gutenkunst, K. S. Brown, C. R. Myers, P. W. Brouwer, V. Elser, and J. P. Sethna, Phys. Rev. Lett. 97, 150601 (2006).

[35] M. K. Transtrum, B. B. Machta, K. S. Brown, B. C. Daniels, C. R. Myers, and J. P. Sethna, J. Chem. Phys. 143 (2015), 10.1063/1.4923066.

[36] S. Hochreiter and J. Schmidhuber, Neural Comput. 9, 1 (1997).

[37] N. S. Keskar, D. Mudigere, J. Nocedal, M. Smelyanskiy, and P. T. P. Tang, arXiv (2016), 10.48550/arXiv.1609.04836, 1609.04836.

[38] Y. Feng and Y. Tu, Proc. Natl. Acad. Sci. U.S.A. 118, e2015617118 (2021).

[39] A. L. Fairhall, G. D. Lewen, W. Bialek, and R. R. d. R. Van Steveninck, Nature 412, 787 (2001), 11518957.

[40] I. Dean, N. S. Harper, and D. McAlpine, Nat. Neurosci. 8, 1684 (2005).

[41] G. A. Jacobson, P. Rupprecht, and R. W. Friedrich, Curr. Biol. 28, 1 (2018).

[42] W.F. Młynarski and A. M. Hermundstad, eLife (2018), 10.7554/eLife.32055.

[43] A. I. Weber, K. Krishnamurthy, and A. L. Fairhall, Annu. Rev. Vision Sci. 5:427–449. (2019), 10.1146/annurev-vision-091718-014818, 31283447.

[44] W.F. Młynarski and A. M. Hermundstad, Nat. Neurosci. 24, 998 (2021).

[45] A. Torralba and A. Oliva, Network 14, 391 (2003).

[46] B. A. Olshausen and D. J. Field, Network 7, 333 (1996).

[47] M. A. Goldin, B. Lefebvre, S. Virgili, M. K. Pham Van Cang, A. Ecker, T. Mora, U. Ferrari, and O. Marre, Nat. Commun. 13, 1 (2022).

[48] C. Chen and W. G. Regehr, Neuron 28, 955 (2000), 11163279.

[49] A. R. Chandrasekaran, R. D. Shah, and M. C. Crair, J. Neurosci. 27, 1746 (2007).

[50] E. Y. Litvina and C. Chen, Neuron 96, 330 (2017).

[51] Y.-t. Li and M. Meister, eLife 10.7554/eLife.82367.

[52] B. D. Hoshal, C. M. Holmes, K. Bojanek, J. M. Salisbury, M. J. Berry, O. Marre, and S. E. Palmer, Proc. Natl. Acad. Sci. U.S.A. 121, e2313676121 (2024).

[53] R. Ramesh, A. Bisulco, R. W. DiTullio, L. Wei, V. Balasubramanian, K. Daniilidis, and P. Chaudhari, arXiv (2024), 10.48550/arXiv.2407.13841, 2407.13841.

[54] R. Linsker, Computer 21, 105 (1988).

[55] R. Linsker, Advances in Neural Information Processing Systems 1 (1988).

[56] T. Sanger, Advances in Neural Information Processing Systems 1 (1988).

[57] T. D. Sanger, Neural Networks 2, 459 (1989).

[58] P. J. B. Hancock, R. J. Baddeley, and L. S. Smith, Network 3, 61 (1992).

[59] B. A. Olshausen and D. J. Field, Nature 381, 607 (1996).

[60] E. Schneidman, M. J. Berry, R. Segev, and W. Bialek, Nature 440, 1007 (2006).

[61] G. Tkacik, E. Schneidman, M. J. Berry Ii, and W. Bialek, arXiv (2006), 10.48550/arXiv.q-bio/0611072, q-bio/0611072.

[62] T. Mora, S. Deny, and O. Marre, Phys. Rev. Lett. 114, 078105 (2015).

[63] G. Tkačik, O. Marre, D. Amodei, E. Schneidman, W. Bialek, and M. J. B. I. I., PLoS Comput. Biol. 10, e1003408 (2014).

[64] G. Tkačik, T. Mora, O. Marre, D. Amodei, S. E. Palmer,M. J. Berry, and W. Bialek, Proc. Natl. Acad. Sci. U.S.A. 112, 11508 (2015).

[65] J. S. Prentice, O. Marre, M. L. Ioffe, A. R. Loback, G. Tkačik, and M. J. B. Nd, PLoS Comput. Biol. 12, e1005148. (2016), 27855154.

[66] A. Loback, J. Prentice, M. Ioffe, and M. Berry Ii, Neural Comput. 29, 3119 (2017).

[67] K. Ruda, J. Zylberberg, and G. D. Field, Nat. Commun. 11, 1 (2020).

[68] L. Ramirez, W. Bialek, S. E. Palmer, and D. J. Schwab, arXiv (2021), 10.48550/arXiv.2112.14334, 2112.14334.

[69] S. Wang, B. Hoshal, E. de Laittre, O. Marre, M. Berry, and S. Palmer, Advances in Neural Information Processing Systems 35, 11355 (2022).

[70] J. Hubbard, Phys. Rev. Lett. 3, 77 (1959).

[71] R. L. Stratonovich, Soviet Physics Doklady 2, 416 (1957).

[72] T. Plefka, J. Phys. A: Math. Gen. 15, 1971 (1982).

[73] T. Tanaka, Neural Comput. 12, 1951 (2000).

[74] M. J. Schnitzer and M. Meister, Neuron 37, 499 (2003).

[75] E. Ganmor, R. Segev, and E. Schneidman, Proc. Natl. Acad. Sci. U.S.A. 108, 9679 (2011).

[76] E. Schneidman, J. L. Puchalla, R. Segev, R. A. Harris, W. Bialek, and M. J. Berry, J. Neurosci. 31, 15732 (2011).

[77] A. Krizhevsky, , 32 (2009).

[78] D. Kunin, J. M. Bloom, A. Goeva, and C. Seed, arXiv (2023), (2019), 10.48550/arXiv.1901.08168, 1901.08168.

[79] X. Bao, J. Lucas, S. Sachdeva, and R. Grosse, arXiv (2020), 10.48550/arXiv.2007.06731, 2007.06731.

[80] D. L. Ruderman and W. Bialek, Phys. Rev. Lett. 73, 814 (1994).

[81] U. Ferrari, C. Gardella, O. Marre, and T. Mora, eNeuro 4 (2017), 10.1523/ENEURO.0166-17.2017.

[82] D. W. Dong and J. J. Atick, Network 6, 345 (1995).

[83] G. Mahuas, T. Buffet, O. Marre, U. Ferrari, and T. Mora, PRX Life 3, 033012 (2025).

[84] T. E. Yerxa, E. Kee, M. R. DeWeese, and E. A. Cooper, PLoS Comput. Biol. 16, e1008146 (2020).

[85] J. Gjorgjieva, M. Meister, and H. Sompolinsky, PLoS Comput. Biol. 15, e1007476 (2019).

[86] J. Sohl-Dickstein, arXiv 10.48550/arXiv.1205.4295, 1205.4295. (2012),

[87] J. Sohl-Dickstein, P. B. Battaglino, and M. R. DeWeese, Phys. Rev. Lett. 107, 220601 (2011).

[88] I. J. Good, Biometrika 40, 237 (1953).

[89] R. Haslinger, D. Ba, R. Galuske, Z. Williams, and G. Pipa, Front. Comput. Neurosci. 7, 44053 (2013).

[90] A. Isihara, J. Phys. A: Gen. Phys. 1, 539 (1968).

[91] B. Wu and J. Pillow, arXiv (2025), 10.48550/arXiv.2512.12467, 2512.12467.

